# Improved anti-tumor vaccination activity of necroptotic cells lacking ATG16L1-dependent unconventional autophagy

**DOI:** 10.64898/2026.10.16.759377

**Authors:** Julia Bandera-Linero, Daniel Oña-Sánchez, Kostantinos Stamatakis, Felipe X. Pimentel-Muiños

## Abstract

Immunogenic forms of tumor cell death can induce effective anti-cancer immune responses, but the molecular mechanisms that regulate the immunogenic potential of cell suicide and how they shape anti-tumor immunity *in vivo* are issues that remain to be fully elucidated. Here we show that absence of unconventional autophagy mediated by the WD40 domain (WDD) of ATG16L1 increases the anti-tumor vaccinating capacity of dying cells. Necroptotic mouse carcinoma MC38 cells lacking the WDD fail to mount a concomitant unconventional autophagic response and exhibit superior vaccinating capacity against MC38 tumors inoculated afterwards. The mutant strain secretes higher levels of ATP during cell demise, whereas other Damage-Associated Molecular Patterns (DAMPs) remain unaffected. The vaccinating effect is accompanied by increased tumor immune infiltration, depends on secreted ATP and requires T cells. These results indicate that suppression of ATG16L1-mediated unconventional autophagy during tumor cell death improves anti-tumor immunity, and identify this process as a potential therapeutic target to enhance the immune contribution to control tumor development.

## Introduction

A diverse array of programmed cell death processes has been described to date, including apoptosis, necroptosis, pyroptosis, ferroptosis as well as a number of other less characterized modalities^1^. In addition to involving different execution mechanisms, these forms of cellular suicide differ in their ability to activate the innate immune system and eventually promote an adaptive immune response directed against the dying cells^2^. For instance, necroptosis is a highly immunogenic cell death routine able to elicit immune activity *in vivo*^3,4,5^. Apoptosis, as the main cell death modality that operates under homeostatic conditions, is typically immunosilent^6^, but immunogenic variants have also been described^7^. The immunogenic potential of cell death is highly relevant in the context of cancer, as immunogenic chemotherapy causes more profound and durable tumor remissions by engaging the immune system to both restrain tumor growth and prevent recurrence through to the development of immune memory^8^.

The onset of immunogenicity during cell death requires the emission of mediators able to stimulate the innate immune properties of the phagocytes that are attracted to the site of cell demise to clear the remains^9^. Such mediators are known as Damage-Associated Molecular Patterns (DAMPs) and include the liberation of ATP, exposure of the endoplasmic reticulum protein calreticulin at the surface of the dying cells, and release of the non-histone nuclear molecule HMGB1 to the extracellular medium^10^. ATP acts in this context both as a find-me factor that recruits phagocytic cells by engaging their purinergic P2Y2 receptors^11^, and also as a pro-inflammatory stimulus via P2X7 receptors for inflammasome activation^12,13^. Calreticulin exposure provides an eat-me signal that promotes phagocytosis of the cellular corpses^14^, whereas secreted HMGB1 stimulates Toll-like receptors (TLRs) and contributes to activate the recruited phagocytes^15^. In addition to these classical DAMPs, cell death is in some cases accompanied by the release of mitochondrial DNA (mtDNA)^16,17^, whose abnormal presence in the cytoplasm stimulates the cGAS/STING signaling system that stimulates production of type I interferons^18,19^. It has been shown that this response can favor the induction of anti-tumor immunity^20,21,22^, although it can also mediate tumor expansion in a context-dependent manner^22,23,24^. Phagocytes activated by these DAMPs trigger a local inflammatory reaction that facilitates their differentiation into efficient antigen presenting cells which, in turn, migrate to the local lymph nodes to promote T cell activation and adaptive immunity^25^.

DAMP emission during the course of cell death requires the induction of a previous stress response that progresses to suicide after insufficient damage resolution^26^, although the detailed mechanisms that regulate the activation of each DAMP in this context are incompletely characterized. Given the importance that involving the immune system has for the therapeutic success of chemotherapy, current efforts to screen for potential chemotherapeutic agents are focusing on their immunogenic capabilities, rather than solely on their plain cytotoxic activity against tumor cells^27^. Early studies identified families of chemotherapeutic compounds particularly prone to inducing immunogenic apoptosis, leading for example to the identification of anthracyclines (in particular mitoxantrone, MTX) as agents holding this property^14,28,29^. In addition to developing increasingly immunogenic drugs, identification of the mechanisms that regulate DAMP exposure could offer opportunities for therapeutic intervention in order to maximize DAMP activation and immunogenic cell death with potential benefits in cancer treatment.

Autophagy, a catabolic pathway that degrades cytoplasmic components^30^ and also plays a role in unconventional secretory processes^31,32^, has been involved in ATP^33,34,35^ and HMGB1^36,37^ liberation during apoptosis, thus contributing to boost the immunogenic potential of this cell death program. In addition to inducing formation of canonical double-membrane autophagosomes, the autophagic machinery participates in unconventional events where single-membrane intracellular structures are labelled with the autophagic marker LC3^38,39,40,41^. ATG16L1, a critical effector that mediates the formation of canonical double-membrane autophagosomes by nucleating an LC3-lipidation complex^30^, is also a central actor in the induction of unconventional autophagic processes^38,39,40,41^. Thus, ATG16L1 responds to upstream activators that engage its C-terminal WD40 domain (WDD; residues 320-607), which is dispensable for canonical autophagy^42,43,44,45^, to induce LC3 lipidation and labeling of single-membrane structures with LC3 without the intervention of the protein complexes that initiate autophagosome formation in the canonical route^46,47,48^.

We previously described that mitochondrial apoptosis is accompanied by an unconventional autophagic response that represses ATP secretion during the course of cell death^49^. This activity requires the ATG16L1 WDD, and correlates with the generation of LC3-positive, single-membrane, ATP-containing vesicles whose inhibition increases ATP liberation^49^. Here we have examined the capacity of unconventional autophagy to control the immunogenic potential of cell death using an *in vivo* experimental system based on the murine tumor cell line MC38. We found that absence of the unconventional autophagic response during MC38 necroptosis increases ATP liberation and enhances the ability of the dying cells to vaccinate against the growth of the same tumor cells inoculated afterwards.

## Results

### Engineering MC38 cells to express full-length or ΔWDD (Nt) forms of ATG16L1

To explore a possible role of unconventional autophagy mediated by the ATG16L1 WDD in the control of the immunogenic potential of tumor cell death *in vivo*, we resorted to the C57BL/6 syngeneic mouse colon adenocarcinoma cell line MC38. We first engineered these cells to express full-length (FL) ATG16L1 or a deleted version lacking the WDD (residues 1-299; Nt) using the CRISPR/Cas9 system. Endogenous ATG16L1 expression was efficiently reduced by using the selected sgRNA (Fig. 1A), and the resulting cells were then restored with sgRNA-resistant retroviral constructs expressing FL or Nt forms of ATG16L1 (Fig. 1A). Both cell lines attained comparable levels of canonical autophagy, as revealed by treatment with bafilomycin (which reveals the basal autophagic flux; Suppl. Figs. 1A and 1C, left and right panels) or rapamycin (a classical activator of the canonical route; Suppl. Figs. 1B and 1C, left and right panels), using in both cases the lipidation of LC3 and its translocation to a vesiculated intracellular pattern as reporter systems to measure the autophagic flux.

**Figure 1.**
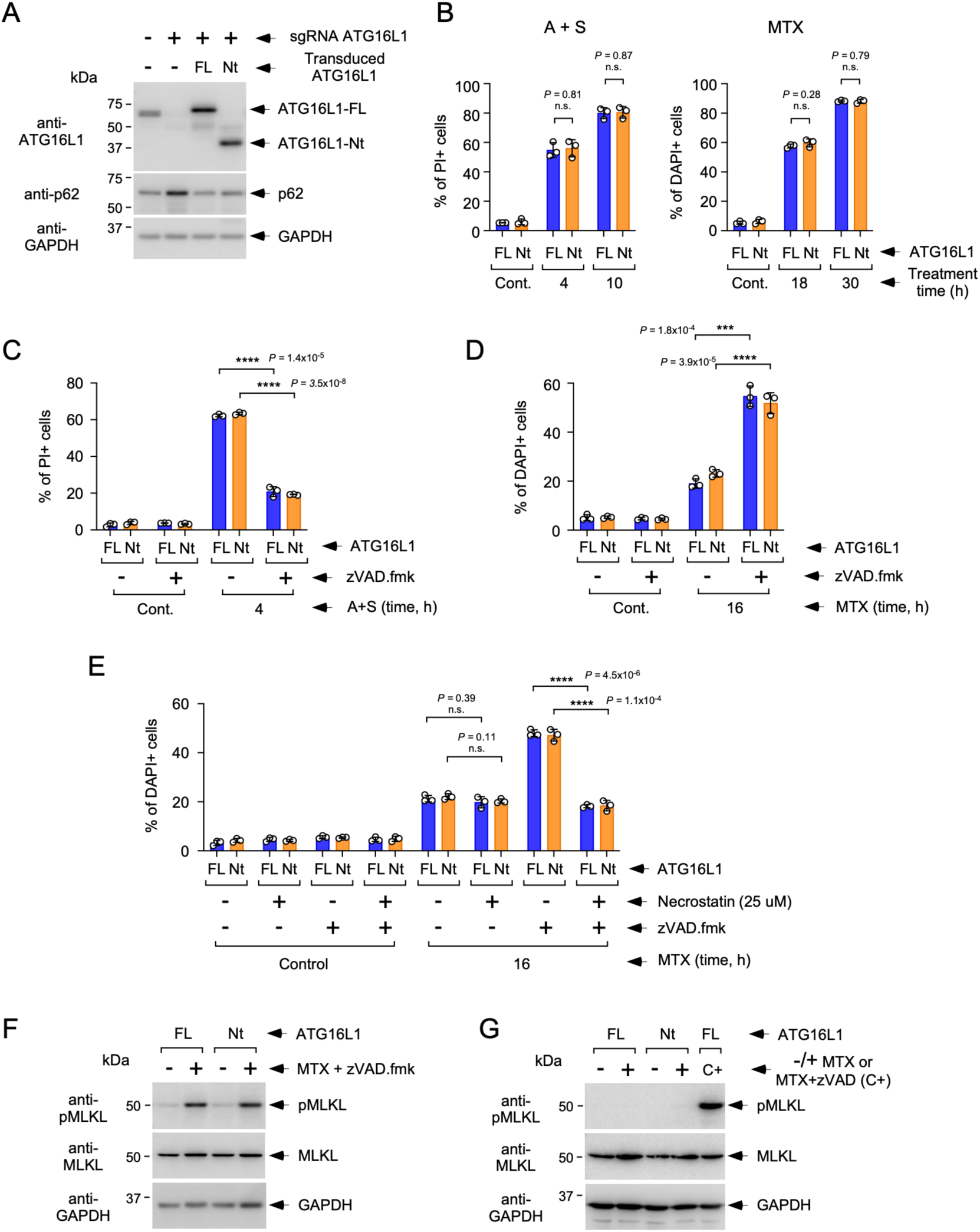
Generation of MC38 strains expressing ATG16L1 full-length (FL) or a truncated version lacking the WDD (residues 1-299; Nt) and characterization of their cell death responses to different stimuli. **(A)** MC38 cells were transduced with a lentivirus expressing a CRISPR/Cas9 sgRNA targeting endogenous ATG16L1 (+) or a control sgRNA (-), and subsequently transduced with retroviruses encoding the indicated versions of ATG16L1 (FL or Nt) or control vector (-). Cells were selected and lysed for Western blot against the indicated molecules. **(B)** The indicated cells (FL or Nt) were treated for the shown times with A+S (5 μM each, left panel), MTX (1.5 μM, right panel) or left untreated (Cont.), and stained with propidium iodide (PI, left) or DAPI (right) to measure cell death by flow cytometry. Shown are mean values -/+ s.d. of the percentage of PI- or DAPI-positive cells obtained from triplicate experimental points (n=3 biological replicas; n.s. *P*>0.05 two-tailed Student’s t test). **(C)** The indicated cells were treated with A+S (5 μM each) in the absence or presence of zVAD.fmk (20 μM) and processed as in B to measure cell death by PI staining. Data are displayed as in B (\*\*\*\**P*<0.001, two-tailed Student’s t test). **(D)** The indicated cells were treated with MTX (1.5 μM) in the absence or presence of zVAD.fmk (20 μM) and processed as in B to measure cell death by DAPI staining. Data are displayed as in B (\*\*\**P*<0.001, \*\*\*\**P*<0.001, two-tailed Student’s t test). **(E)** Cells were treated with MTX (1.5 μM) in the absence or presence of zVAD.fmk (20 μM) or Necrostatin (25 μM) and processed as in B to measure cell death by DAPI staining. Data are displayed as in B (n.s. *P*>0.05, \*\*\*\**P*<0.001, two-tailed Student’s t test). **(F)** Cells were treated with MTX (1.5 μM) and zVAD.fmk (20 μM) for 16 h and lysed for Western blotting against the indicated molecules. **(G)** Cells were treated with MTX (1.5 μM) for 19 h and lysed for Western blotting against the indicated molecules. A positive control sample (C+) from FL cells treated with MTX (1.5 μM) and zVAD.fmk (20 μM) for 16 h was included in the last lane.

### Cell death responses of engineered MC38 cells

As a previous step to characterize the immunogenic properties of FL and Nt MC38 cells during programmed cell death, we first studied their death responses to a variety of stimuli. We found that the kinetics of cell demise was comparable between FL and Nt MC38 strains in all cases, including the simultaneous treatment with the BH3 mimetics ABT-737 and S63845 (A+S) that activate the intrinsic apoptotic pathway (Fig. 1B, left panel), the established immunogenic chemotherapy mitoxantrone (MTX; Fig. 1B, right panel) and other chemotherapy agents like cisplatin and 5-fluorouracil (5-FU; Suppl. Fig. 2A). Consistent with a predominantly apoptotic mechanism of action, cell death induced by A+S was inhibited by the pan-caspase inhibitor zVAD.fmk (Fig. 1C). However, unexpectedly, zVAD.fmk robustly enhanced MTX-induced cell death, again irrespective of the ATG16L1 version expressed by the cells (Fig. 1D). Caspase inhibition is required for the induction of necroptosis, a caspase-independent form of programmed cell death that involves plasma membrane poration caused by oligomers of the pseudokinase MLKL, which are assembled in response to RIP kinase family members^5,50^. Treatment with the RIPK1 inhibitor necrostatin reduced the extent of cell death caused by MTX+zVAD.fmk (MTX+zVAD; Fig. 1E), an inhibitory effect that was quantitatively similar to the one observed in response to TNF, cycloheximide (CHX) and zVAD.fmk (the main experimental paradigm of necroptotic cell death^51,52^; Suppl. Fig. 2B). Surprisingly, the death process induced by MTX remained insensitive to necrostatin treatment (Fig. 1E), suggesting that it does not proceed through a necroptotic pathway. MTX+zVAD-induced cell death was accompanied by phosphorylation of MLKL in both FL and Nt cells (Fig. 1F), whereas the process induced by MTX alone was not (Fig. 1G). These results are consistent with a role of necroptosis in cell death induced by MTX+zVAD but not in response to MTX alone.

To explore with further detail the nature of MTX-induced cell death in this system, we measured other parameters of apoptosis. MTX caused caspase-3 processing (Fig. 2A) and release of cytochrome c to the cytoplasm (Fig. 2B), both events linked to the activation of the intrinsic apoptotic pathway that relies on the function of Bcl-2-family members^53,54^. Consistent with an involvement of this pathway, depletion of BAK and BAX (Fig. 2C) prevented cytochrome c mobilization (Fig. 2D) caused by MTX. Surprisingly, MTX+zVAD-induced cell death was also accompanied by cytochrome c release (Fig. 2E), which was BAK,BAX dependent in both FL and Nt cells (Fig. 2F). Cell death induced by MTX was at least partially inhibited by the depletion of BAK and BAX (Fig. 2G), whereas the process triggered by MTX+zVAD was completely insensitive (Fig. 2H).

**Figure 2.**
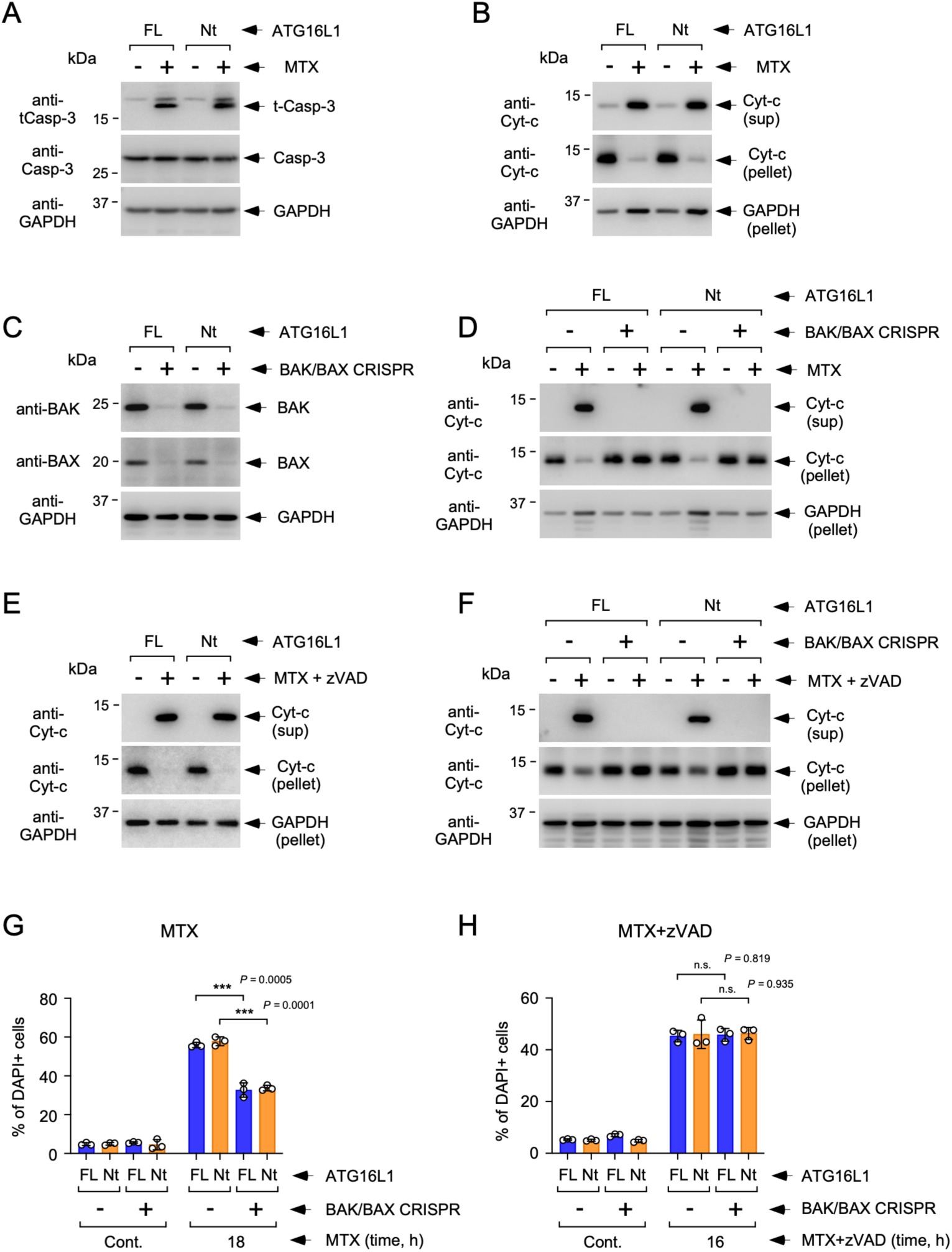
Characterization of cell death induced by MTX and MTX+zVAD.fmk in MC38 cells expressing ATG16L1 FL or Nt. **(A)** Cells expressing the indicated forms of ATG16L (FL or Nt) were treated with MTX (1.5 μM, 19 h) and lysed for Western blot against the shown molecules. **(B)** Cells were treated with MTX (1.5 μM, 14 h) and lysed for evaluation of cytochrome c (Cyt-c) release by Western blotting of protein extracts obtained from the soluble (sup) and insoluble (pellet) fractions. **(C)** Cells were transduced with CRISPR/Cas9 lentiviral particles expressing a control sgRNA species (-) or a mix of particles expressing sgRNAs targeting BAK and BAX (+), and lysed for Western blot using the indicated antibodies. **(D)** The indicated cells were treated with MTX (1.5 μM, 14 h) and lysed to evaluate Cyt-c release as in B. **(E)** Cells were treated with MTX (1.5 μM) + zVAD (20 μM) for 12 h and lysed to evaluate Cyt-c release as in B. **(F)** The indicated cells were treated with MTX (1.5 μM) + zVAD (20 μM) for 12 h and lysed to evaluate Cyt-c release as in B. **(G)** The indicated cells were treated with MTX (1.5 μM) for 18 h and processed to measure cell death by DAPI staining. Shown are mean values -/+ s.d. of the percentage of DAPI-positive cells obtained from triplicate experimental points (n=3 biological replicas; *** *P*<0.001, two-tailed Student’s t test). **(H)** The indicated cells were treated with MTX (1.5 μM) + zVAD.fmk (20 μM) for 16 h and processed to measure cell death by DAPI staining. Shown are mean values -/+ s.d. of the percentage of DAPI-positive cells obtained from triplicate experimental points (n=3 biological replicas; n.s. *P*>0.05, two-tailed Student’s t test).

These data argue that MTX kills MC38 cells by activating the intrinsic mitochondrial pathway to apoptosis mediated by cytochrome c release and caspase activation. However, this route is efficiently redirected to a necroptotic cell death modality by caspase inhibition (MTX+zVAD in this context) while still conserving intrinsic apoptotic features that do not appear to participate in the death execution process, like cytochrome c mobilization. Notably, FL and Nt cells behaved similarly in all these assays, indicating that absence of the WDD does not influence the course of a variety of cell death modalities. Since necroptosis has been described as a particularly immunogenic form of programmed cell death^3,4,50^ we were interested in studying whether unconventional autophagy is activated in response to MTX+zVAD to control the immunogenic properties of necroptosis induced in MC38 cells.

### Unconventional autophagy is induced in response to various cell death stimuli

Treatment with MTX+zVAD provoked an LC3 lipidation process in FL MC38 cells that was reduced in the Nt strain (Fig. 3A, left panel), suggesting the involvement of unconventional WDD-dependent autophagic mechanisms. Consistently, a similar result was obtained in the presence of a VPS34 inhibitor (VPS34-INH; Fig. 3A, right panel) that is able to block canonical autophagy, as shown in control assays carried out in response to bafilomycin (Suppl. Fig. 2C). These results were recapitulated by measuring translocation of GFP-LC3 to a vesiculated pattern (Fig. 3B, top and bottom panels). Unconventional autophagic mechanisms mediated by ATG16L1 WDD induce the Conjugation of ATG8 to Single Membranes (CASM) in response to lysosome alterations initially sensed by the vesicular ATPase (vATPase)^40^, a process that is inhibited by the *Salmonella* virulence factor SopF^55^. Expression of GFP-SopF robustly inhibited LC3 activation caused by MTX+zVAD in the presence of VPS34-INH (Fig. 3C), pointing to the idea that CASM is likely the main mechanism driving LC3 lipidation in this experimental paradigm. WDD-dependent, VPS34-INH-resistant LC3 lipidation and translocation to a punctated pattern was also detected in response to A+S (Figs. 3D and 3E, top and bottom panels) and MTX (Figs. 3F and 3G, top and bottom panels), as well as other chemotherapy agents (Suppl. Fig. 2D), indicating that induction of unconventional autophagy is a common event present during a variety of cell death routines. Notably, depletion of both BAK and BAX prevented LC3 lipidation caused by MTX+zVAD (Suppl. Fig. 2E; left panel) and MTX (Suppl. Fig. 2E; right panel), pointing to the idea that the autophagic response requires engagement of the mitochondrial apoptotic machinery.

**Figure 3.**
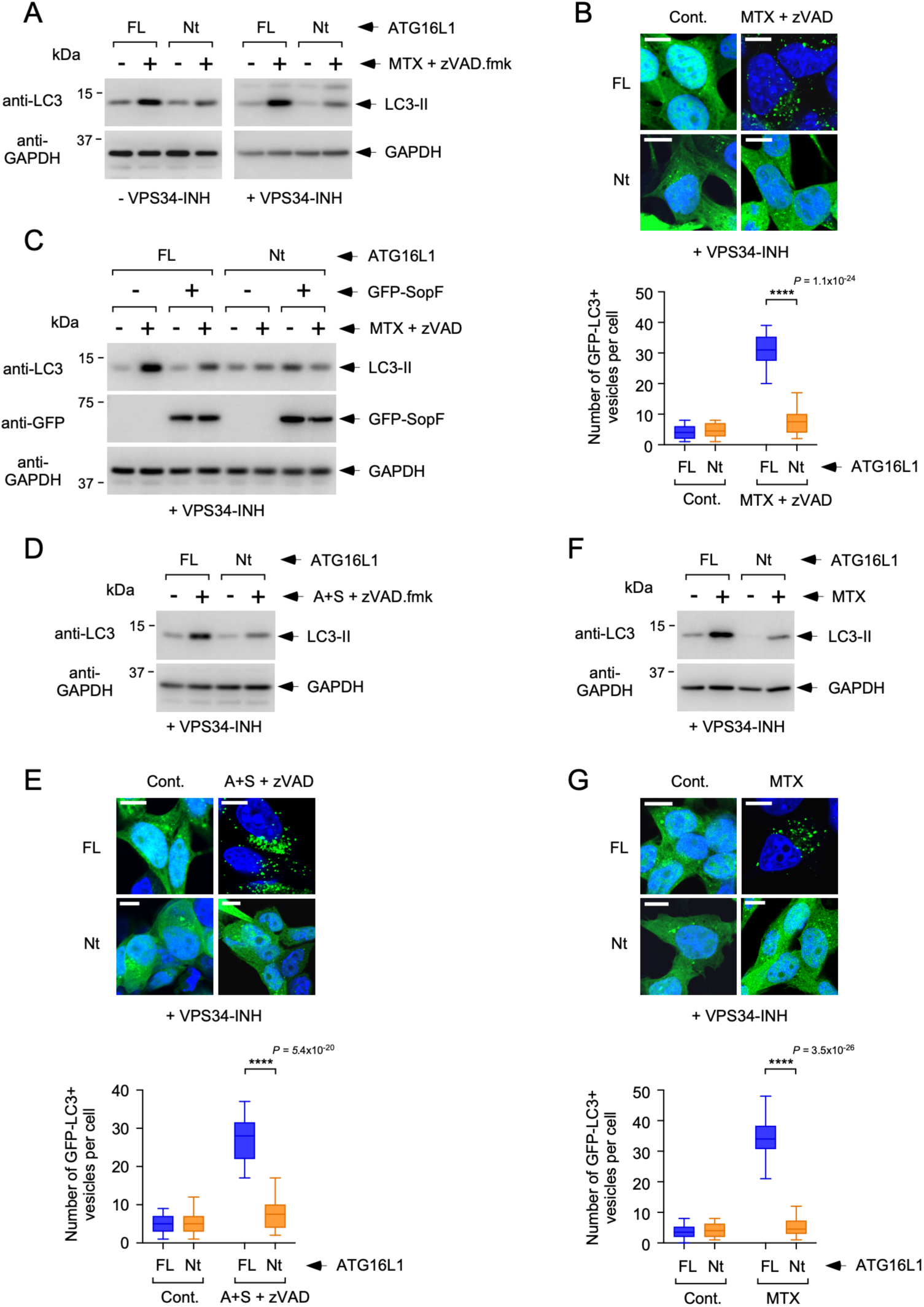
Unconventional autophagy induced during cell death induced by different stimuli. **(A)** The indicated cells (FL or Nt) were treated with MTX (1.5 μM) + zVAD (20 μM) for 16 h in the absence or presence of 2 μM of the VPS34 inhibitor (VPS34-INH, as displayed) and lysed for Western blot against the indicated molecules. **(B)** The indicated cells stably expressing GFP-LC3 were treated for 16 h with MTX (1.5 μM) + zVAD (20 μM) in the presence of 2 μM of the VPS34 inhibitor and fixed for microscopy. Shown are representative confocal pictures (top panel) and quantification of the number of GFP-LC3-puncta per cell (bottom panel). Data are displayed as box-plots where the central line indicates the median value, the box shows percentiles 25-75 and the whiskers include the most extreme values (n=26 cells; \*\*\*\**P*<0.0001 two-tailed Student’s t test). **(C)** The indicated cells expressing (+) or not (-) GFP-SopF via retroviral transduction were treated with MTX (1.5 μM) + zVAD (20 μM) for 16 h in the presence of VPS34-INH (2 μM) and lysed for Western blot against the indicated molecules. **(D)** Cells were treated with A+S (5 μM each) for 8 h in the presence of zVAD (20 μM) and VPS34-INH (2 μM) and lysed for Western blot against the indicated molecules. **(E)** The indicated cells stably expressing GFP-LC3 were treated with A+S (5 μM each) for 6 h in the presence of zVAD (20 μM, to prevent cell detachment and death) and VPS34-INH (2 μM) and fixed for confocal microscopy. Representative confocal pictures (top panel) and quantification of the number of GFP-LC3-puncta per cell (bottom panel) are displayed as in B. **(F)** Cells were treated with MTX (1.5 μM) for 19 h in the presence of VPS34-INH (2 μM) and lysed for Western blot against the indicated molecules. **(G)** Cells stably expressing GFP-LC3 were treated with MTX (1.5 μM) for 18 h in the presence of VPS34-INH (2 μM) and fixed for confocal microscopy. Representative confocal pictures (top panel) and quantification of the number of GFP-LC3-puncta per cell (bottom panel) are displayed as in B.

### Unconventional autophagy suppresses ATP mobilization during cell death induced by MTX+zVAD

To establish the relevance of the noncanonical autophagic route in the control of immunogenic cell death, we measured the mobilization of different DAMPs in FL and Nt cells in response to the different stimuli. Treatment with MTX+zVAD caused increased ATP secretion in cells lacking the WDD (Fig. 4A; left panel), indicating that our previous results showing that unconventional autophagy represses ATP secretion during apoptosis^49^ can unexpectedly be extended to necroptotic cell death. In fact, we also detected high levels of ATP trapped into the LC3-positive unconventional vesicles generated in FL cells, as established by their staining with the ATP-sensitive reagents quinacrine (Suppl. Fig. 3A) and Bodipy-ATP (Suppl. Fig 3B). MTX also caused significantly enhanced ATP secretion in Nt cells, although the effect was rather discrete (Fig. 4A; middle panel), whereas A+S induced equal ATP release in both cell lines (Fig. 4A; right panel). In addition, both MTX+zVAD and MTX alone provoked HMGB1 secretion to a comparable extent in FL and Nt cells (Fig. 4B; left and middle panels, respectively), while A+S barely induced any secretion at all in both cell lines (Fig. 4B; right panel). Treatment with MTX+zVAD induced calreticulin exposure at the surface of the dying cells (Fig. 4C; left panel) which was slightly higher in cells treated with A+S (Fig. 4C; right panel), although no differences between FL and Nt cells were detected. On the contrary, MTX caused poor calreticulin exposure in both cell lines (Fig. 4C; middle panel). While highly context-dependent, STING activation during cell death can contribute to boost the immunogenicity of cell suicide proceses^20,21,22^. However, STING signaling is known to be inhibited by caspases^56^, so we measured STING phosphorylation under the death-inducing conditions that included treatment with the pan-caspase inhibitor zVAD.fmk. MTX+zVAD was unable to trigger STING phosphorylation in any of the cell lines (Fig. 4D, left panel), whereas the STING agonist dimethylxanthenone acetic acid (DMXAA; DMX) caused robust activation in both of them (Fig. 4D, right panel). The latter result shows that the STING signaling system is fully functional in MC38 cells, indicating a deficient activation specifically during cell death induced by MTX+zVAD.

**Figure 4.**
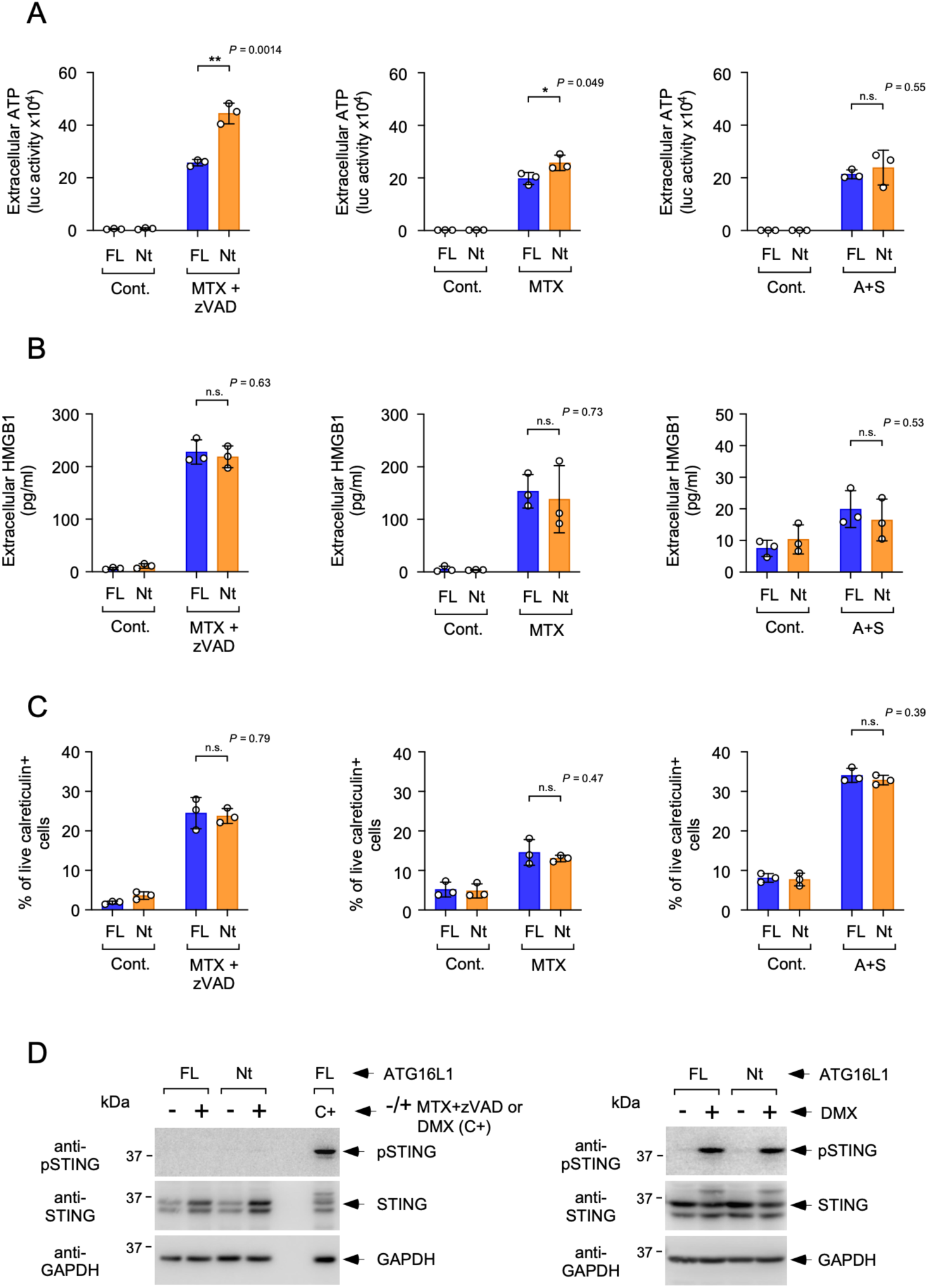
DAMP emission during cell death induced by different stimuli. **(A)** The indicated cellular strains were treated as indicated with MTX+zVAD (1.5 μM and 20 μM, respectively; left panel), MTX (1.5 μM; middle panel) or A+S (5 μM each; right panel) and the levels of extracellular ATP were measured at 14 and 20 h (for MTX+zVAD), 16 and 22 h (for MTX) or 4 and 9 h (for A+S) post-treatment. The graphs show mean values-/+ s.d. of cumulative data resulting from the sum of luciferase activity units obtained at both time points from triplicate experimental points (n=3 biological replicas; n.s. *P*>0.05, \**P*<0.05, \*\**P*<0.01 two-tailed Student’s t test). **(B)** The indicated cells were treated as indicated with MTX+zVAD (1.5 μM and 20 μM, respectively, 20 h; left panel), MTX (1.5 μM, 24 h; middle panel) or A+S (5 μM each, 12 h; right panel) and the levels of extracellular HMGB1 were measured by ELISA. The graphs show mean values -/+ s.d. of HMGB1 levels (pg/ml) obtained from triplicate experimental points (n=3 biological replicas; n.s. *P*>0.05, two-tailed Student’s t test). **(C)** The indicated cells were treated as indicated with MTX+zVAD (1.5 μM and 20 μM, respectively, 16 h; left panel), MTX (1.5 μM, 20 h; middle panel) or A+S (5 μM each, 6 h; right panel) and the surface expression of calreticulin was evaluated by flow cytometry in the live cell compartment. The graphs show mean values -/+ s.d. of the percentage of live cells expressing CD39 obtained from triplicate experimental points (n=3 biological replicas; n.s. *P*>0.05, two-tailed Student’s t test). **(D)** Cells were treated with MTX (1.5 μM) + zVAD.fmk (20 μM) for 16 h (left panel) or with DMXAA (DMX, 25 μg/ml) for 2 h (right panel), and lysed for Western blotting against the indicated molecules. A positive control sample (C+) obtained from FL cells treated with DMXAA (DMX, 25 μg/ml, 2 h) was included in the last lane of the experiment displayed on the left panel.

Together, these studies establish that MTX+zVAD causes robust exposure of most tested DAMPs in MC38 cells, and also show that ATP liberation is negatively regulated by the unconventional autophagic response that is induced in parallel during the suicide process induced by these stimuli. Given the relevance that ATP liberation has in boosting the immunogenic properties of programmed cell death^33,35^, these data suggest that inhibition of such unconventional autophagic pathway might be able to enhance the immunogenicity of necroptotic MC38 cells treated with MTX+zVAD.

### Necroptotic MC38-Nt cells show increased anti-tumor immunogenic activity *in vivo*

To explore the role of the unconventional autophagic response in the control of anti-tumor immunogenicity *in vivo* we resorted to vaccination assays where FL and Nt cells dying in response to MTX+zVAD were inoculated subcutaneously in the flank of syngeneic C57BL/6 mice. One week later, unaltered MC38 cells constitutively expressing the luciferase gene were injected in the contralateral flank and tumor growth was subsequently followed over time. Initial studies showed that vaccination with FL cells caused the tumors to grow slower compared to unvaccinated control mice (Fig. 5A). Interestingly, vaccination with Nt cells appeared to have an increased ability to do so (Fig. 5A), although in this case the difference was not statistically significant. Assays carried out with a higher statistical power (higher n) and specifically designed to measure the difference between FL and Nt vaccination confirmed that MC38 tumors are smaller in mice vaccinated with Nt cells, both by measuring their volume over time (Fig. 5B, left and right panels) and by establishing their luciferase activity *in vivo* (Fig. 5C, left and right panels). Notably, expression of the ecto-ATPase CD39^33,57^ in Nt cells via retroviral transduction (Suppl. Figs. 4A and 4B) was able to reduce the levels of extracellular ATP detected during cell death (Suppl. Figs. 4C) and completely reverted to FL levels the enhanced vaccination effect provided by the mutant strain (Fig. 5D). This result points to the idea that the increased ATP secretion observed in MC38 Nt cells treated with MTX+zVAD at least partially contributes to their increased vaccinating potential *in vivo*. In addition, depletion of T cells via repeated injection of an anti-Thy1 antibody (Suppl. Fig. 4D) abrogated the advantageous effect conferred by absence of the WDD in the vaccinating cells (Fig. 5E), suggesting that vaccination with Nt cells reduces tumor growth by improving an anti-tumor immune response mediated by T cells. Together, these results indicate that absence of the unconventional autophagic response mediated by the WDD improves the immunogenic activity of necroptotic MC38 cells, likely by favoring ATP secretion and the onset of a T cell-dependent immune response.

**Figure 5.**
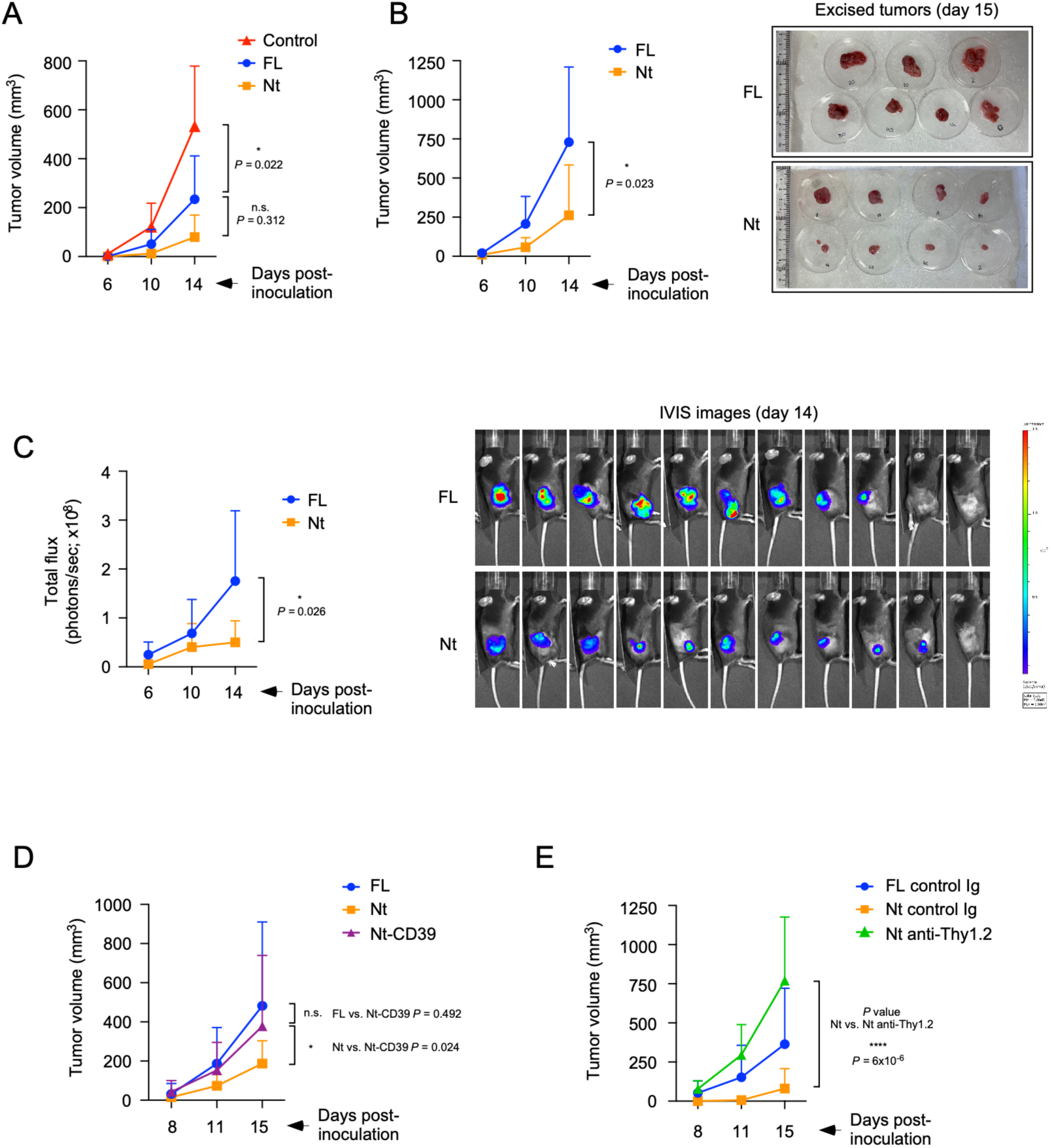
MC38 tumor growth after vaccination with necroptotic MC38 cells expressing ATG16L1 FL or Nt. **(A)** Mice were vaccinated with the indicated MC38 cellular strains treated with MTX (1.5 μM) + zVAD (20 μM) for 15 h (Control: PBS injection), and challenged with wild-type, untreated MC38 cells one week later. The graph shows the evolution of tumor volume measured at the indicated time points after challenge (n=5 mice for PBS, n=6 mice for FL and Nt; n.s. *P*>0.05, \**P*<0.05; Tukey-corrected two-way ANOVA). **(B)** Mice were vaccinated as in A with the indicated dying cells, and challenged also as in A. The graph (left panel) shows the progress of tumor volume measured at the indicated post-challenge time points (n=9 mice; \**P*<0.05; Tukey-corrected two-way ANOVA). The right panel shows pictures of the tumors taken immediately after dissection (day 15 post-challenge). **(C)** Mice were vaccinated and challenged as in B. The graph on the left shows the evolution of tumor size measured as the maximum absolute number of photons emitted per second after D-luciferin injection (total flux) at the indicated post-challenge time points (n=11 mice; \**P*<0.05; Tukey-corrected two-way ANOVA). The right panel shows IVIS pictures of the tumor-bearing mice taken at the maximal luciferase activity obtained after D-luciferin injection. **(D)** Mice were vaccinated as in A with the indicated dying cells (FL, Nt or Nt cells expressing CD39, Nt-CD39) and subsequently challenged. The graph displays the progress of tumor volume measured at the indicated time points after challenge (n=22 mice for FL and Nt-CD39, n=21 mice for Nt; n.s. *P*>0.05, \**P*<0.05; Tukey-corrected two-way ANOVA). **(E)** Mice were vaccinated as in A and subsequently challenged. The shown antibodies (control Ig or anti-Thy1.2) were then repeatedly injected intraperitoneally to deplete T cells. Tumor progress was followed by measuring tumor volume at the indicated post-challenge time points (n=8; \*\*\*\**P*<0.0001; Tukey-corrected two-way ANOVA). T cell depletion was confirmed by flow cytometry (Suppl. Fig. 4D).

### Vaccination with necroptotic ATG16L1-Nt cells increases the immune infiltration of MC38 tumors

To explore with more detail the cellular bases of the improved vaccination effect provided by cells lacking unconventional autophagy, we studied the profile of the immune cells that infiltrate the tumors in the vaccinated mice. Tumors growing in mice inoculated with dying Nt cells contained an increased percentage of the CD45+ immune population compared to those vaccinated with FL controls (Fig. 6A, left panel) and a parallel reduced percentage of CD45-negative cells (Fig. 6A, right panel). When evaluated in the context of the whole tumor, the immune populations that were significantly elevated in mice vaccinated with Nt cells were CD3+ cells as a whole, CD3+CD4+ double-positive T cells, CD11b+F4/80+ cells (macrophages) and CD11b+CD11c+ (dendritic cells) (Fig. 6B), whereas NK cells (CD3-NK1.1+) and CD3+CD8+ T cells remained unchanged (Fig 6B). Consistent with previous studies^58,59^, myeloid cells accounted for the majority of the immune cell infiltration in the tumor mass (macrophages and dendritic cells; Fig. 6B). When corrected against the number of CD45-negative cells (mostly tumor cells), all immune populations showed a tendency to be enriched in those tumors developed in mice vaccinated with Nt cells (Fig. 6C), likely because there is a substantial reduction in tumor cellularity in these conditions (tumor size; see Fig. 5B). Such increased percentages were again significant for CD3+ and CD4+ T cells, macrophages and dendritic cells (Fig. 6C), and also for NK cells in this case (Fig. 6C). On the contrary, the change observed in CD8+ T cells remained below significance (Fig. 6C). To explore the activation status of the infiltrating T cells we measured the expression levels of the surface markers CD25 and CD69. CD4+ T cells expressing both markers (CD3+CD4+CD25+CD69+) were significantly elevated in mice vaccinated with Nt cells, both when measured in the whole tumor (Fig. 6D) and, more prominently, after correction against CD45-negative cells (Fig. 6E). In contrast, the levels of activated CD8+ T cells (CD3+CD8+CD25+CD69+) did not change (Fig. 6D and 6E). Taken together, these results are consistent with the idea that absence of unconventional autophagy in the vaccinating cells strengthens the immune infiltration of MC38 tumors, and point to a particularly relevant role of the infiltrating activated CD4+ T cell compartment in the improved tumor control observed in these conditions.

**Figure 6.**
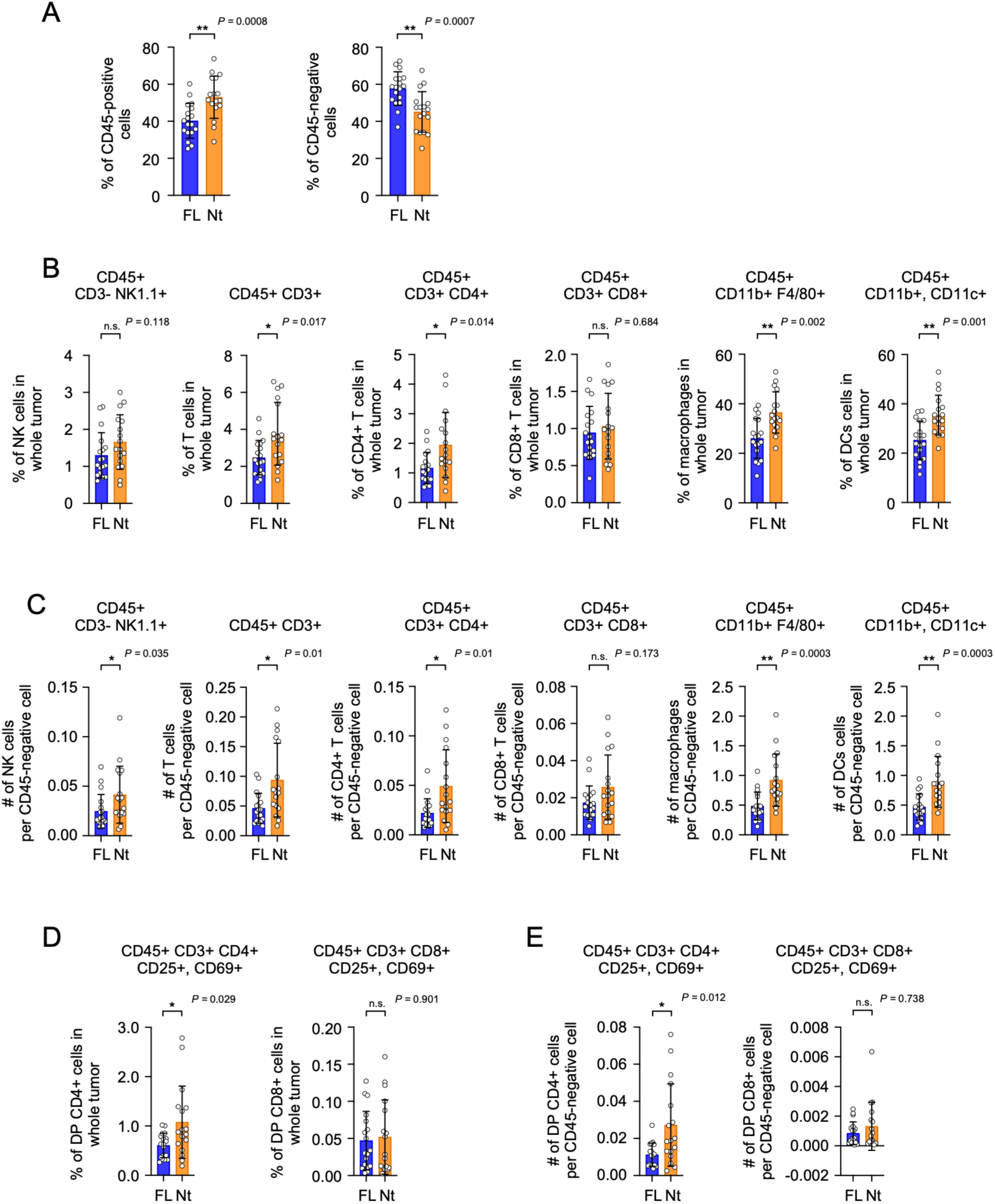
Infiltrating immune profile of subcutaneous MC38 tumors generated after vaccination with dying MC38 FL or Nt cells. (A-E) Mice were vaccinated with the indicated MC38 cells treated with MTX (1.5 μM) + zVAD (20 μM) for 15 h, and subsequently challenged with wild-type, untreated MC38 cells one week later. Tumors were dissected 15 days post-challenge and disaggregated for analysis of the indicated cell populations by flow cytometry. The graphs show mean values -/+ s.d. of the percentage of positive cells in the whole tumor (considering the whole tumor as the sum of the total numbers of CD45+ and CD45-cells; A, B and D), or the number of positive cells divided by the number of CD45-negative cells (mostly tumor cells; C and E) (n=18 mice; n.s. *P*>0.05, \**P*<0.05, \*\**P*<0.01; unpaired nonparametric Mann-Whitney t test).

## Discussion

Here we show that inhibition of an ATG16L-dependent, WDD-mediated unconventional autophagic response that accompanies necroptosis favors the ability of this death program to activate anti-tumor immunity *in vivo*. Such beneficial effect appears to be mediated by the increased secretion of ATP that occurs concomitant to the execution of cell death in WDD-deficient cells. These results are in line with our previous data showing that inhibition of a similar unconventional autophagic phenomenon excited during apoptosis causes enhanced ATP secretion^49^. Interestingly, we found that the onset of unconventional autophagy is a common feature conserved across a variety of cell death routines, raising the notion that it may constitute a general mechanism that controls the immunogenicity of programmed cell death.

Our data show that absence of the WDD does not alter the exposure of any other DAMP aside of ATP during necroptosis induced by MTX+zVAD, and demonstrate that expression of the ecto-ATPase CD39 reverts the vaccination advantage provided *in vivo* by Nt cells. Both results point to a relevant role of ATP in the increased immunogenic potential shown by this strain, an importance that has been raised in other studies^13,33^. However, it remains possible that other unknown DAMPs might also be regulated by noncanonical autophagy and actually contribute to the improved vaccination ability of Nt cells, provided that their action is hierarchically dependent on ATP liberation. In this scenario, if the effect of ATP secretion is neutralized (for example by CD39 expression, as we show here) the impact of such DAMPs would remain undetected. In this regard it is worth mentioning that ATP acts as a find-me factor for phagocyte recruitment^11^, so in the absence of ATP liberation phagocyte recruitment would be defective and any secondary DAMPs would be unable to provide an immunoregulatory effect.

Our data support the idea that absence of unconventional autophagy in the vaccinating cells strengthens the immune infiltration of MC38 tumors. In addition, we observed that the improved anti-tumor response in these conditions requires T cells, since the effect disappears when this immune population is depleted. However, only CD4+ T cells and, in particular, activated CD4+ cells, are increasingly infiltrated in the smaller tumors developed in mice vaccinated with dying Nt cells, whereas the classical cytotoxic CD8+ T cell population remains unchanged. As the immune response to subcutaneous MC38 tumors has been shown to be mainly mediated by CD8+ cytolytic T cells^60^, our results suggest that the increased number of infiltrating activated CD4-positive T cells could be providing improved help to specific CD8+ cytotoxic effectors. Alternatively, these results could indicate that the main effector population in this context are CD4-positive cytotoxic cells, a subset of the T cell compartment that seems to have a relevant role in anti-tumor immunity^61,62^.

Exactly how the unconventional autophagic response controls ATP liberation is currently unclear. Our previous data showed a strong correlation between ATP entrapment inside the single-membrane, LC3-labelled vesicles generated during apoptosis and reduced ATP secretion^49^, and here we find a similar correlation in a necroptotic experimental system. These data suggest that ATP sequestration inside these vacuoles causes reduced ATP release along the death routine. However, it seems unlikely that the observed vesicles may have the capacity to sequester enough ATP to significantly reduce the cytoplasmic pool available for secretion. We believe it is more plausible that the LC3-positive vesicles are sequestering a particular pool of ATP that is destined for secretion, pointing to a mechanism where cytoplasmic ATP is not indiscriminately liberated, but subjected to compartmentalization for a regulated release. However, while this is conceivable in apoptosis, which does not involve membrane permeabilization until the later stages, it is more difficult to imagine in necroptosis, a form of cell death that proceeds with formation of plasma membrane pores that allow the release of cytoplasmic metabolites. Exactly how ATP compartmentalization might be involved in the control of ATP liberation during necroptosis remains to be explored.

Our earlier studies showed that unconventional LC3-positive vesicles are formed in response to BH3-only proteins by a mitochondrial pathway that emanates from active BAK and involves prohibitins and ATG16L1^49^. Being apoptosis and necroptosis such disparate death programs we initially judged as unlikely that there could be a common signaling route that induces the unconventional formation of LC3-positive vesicles. Surprisingly, however, while the death execution modality induced by MTX+zVAD in MC38 cells is necroptotic, key features of mitochondrial apoptosis remain as bystander events, since the death process is accompanied by BAK,BAX-dependent cytochrome c release. Thus, there is a mix of molecular features characteristic of necroptosis and apoptosis, as it has been observed in other experimental systems^63,64^. In this scenario, it is plausible that the BAK-dependent atypical autophagic route that we previously described in an apoptotic system may also be functional during MTX+zVAD-induced necroptosis, an idea supported by our data showing that BAK and BAX are required for the LC3 response in this case. According to these considerations, the LC3-positive atypical autophagic vesicles are probably induced by a similar pathway in both death paradigms. To what extent caspase inhibition can redirect other chemotherapy-induced apoptotic programs to necroptosis while preserving features of mitochondrial apoptosis is presently unclear, but this may be a more general phenomenon.

Previous publications indicate that mitochondrial permeabilization promotes the release of mtDNA, whose presence in the cytoplasm activates the innate immune receptor STING^16,17,18^. Other studies show that STING activation induces the formation of single-membrane, LC3-positive vesicles through an ATG16L1 WDD-dependent mechanism^65^ that requires the channel activity of STING^66^ and is inhibited by SopF^65,66^. Therefore, it is tempting to speculate that the unconventional autophagic pathway we describe here could be mediated by STING. However, we found no signs of STING activation induced by MTX+zVAD, despite the fact that the cGAS/STING system responds robustly to DMXAA. These results indicate that STING is unlikely to be involved in the unconventional response that occurs during necroptosis in the experimental system we describe here. What alternative mechanisms could be involved is currently unclear.

Finally, our data are consistent with a superior ability of necroptosis to elicit anti-tumor immune responses, as it has been proposed previousy^3,4^. More generally, our results suggest that targeted inhibition of WDD-mediated function could constitute an effective therapeutic strategy to increase the immunostimulatory properties of tumor cell death. In fact, since the WDD is dispensable for canonical autophagy, the noncanonical response could conceivably be inhibited with therapeutic purposes without altering the canonical mechanism and its important homeostatic functions, thus helping minimize unwanted side effects.

## Methods

### Cell lines and reagents

MC38 cells were purchased from Kerafast (Ref. ENH204-FP). Cells were cultured at 37°C and a humidified 5% CO_2_ atmosphere, in DMEM (Invitrogen, 52100-047) containing 10% heat-inactivated FBS (Invitrogen, 10437028), L-glutamine (2 mM, TCI, G0063) and 100 U/ml of penicillin/streptomycin (Invitrogen, 15140130). Cells have routinely been tested for mycoplasma contamination.

### Chemicals

zVAD.fmk (Becton Dickinson, 550377), bafilomycin A1 (Sigma, B1793) E64d (EST, Calbiochem, 330005), rapamycin (Selleckchem, S1039), mitoxantrone (MTX, Sigma, M6545), ABT-737 (MedChemExpress, HY-50907), S63845 (MedChemExpress, HY-100741), necrostatin (MedChemExpress, HY-15760), cisplatin (Sigma, 232120), 5-FU (Sigma, 343922), TNF (Peprotech, 315-01A), cycloheximide (Sigma, C7698), VPS34-inhibitor (Selleckchem, Compound 19, S8456), DMXAA (MedChemExpress, HY-10964), DAPI (Sigma, 10236276001), ARL67156 (Sigma, A265), polybrene (Sigma, H9268), quinacrine (Sigma, Q3251), Bodipy-ATP (Thermo Fisher, A12410), puromycin dihydrochloride (Sigma, P8833), hygromycin B (Thermo Fisher, 10687010), blasticidin (Invivogen, ant-bl-1).

### CRISPR/Cas9 and DNA constructs

To deplete protein expression in MC38 cells we used a lentiviral CRISPR/Cas9 system based on the lentiCRISPRv2 vector (which includes a puromycin-selection cassette; Dr. Feng Zhang, Addgene, Ref. 52961; for ATG16L1) or a derivative construct including a blasticidin selection module (Dr. Brett Stringer, Addgene, Ref. 98293; for BAK and BAX). The sgRNA guides were selected using the BreakingCas server (Centro Nacional de Biotecnologia, Madrid, Spain; https://bioinfogp.cnb.csic.es/tools/breakingcas/). Their sequences are: ATG16L1 5’ ctggaagcgtcacatcgcgg 3’; BAK 5’ acaaggaccaggtcccccga 3’ and BAX 5’ gcttcttggtggacgcatcc 3’. Lentiviral plasmids containing the specific guides were co-transfected with the helper plasmids psPAX2 (Addgene Ref. 12260) and pCMV-VSV-G (Addgene, Ref. 8454) into HEK293T cells. Virus-enriched supernatants were harvested 48 h later, filtered against 0.45 μM, and diluted 1:1 in fresh medium containing polybrene (8 μg/ml). For depletion of BAK and BAX, supernatants containing both lentiviruses were mixed 1:1 for co-infection. Infections were carried out by spinning the resulting mix onto the target cells for 1 h (2000 rpm, 32°C). Control cells were transduced with a plasmid carrying an sgRNA species with no known target in mouse cells. Cells were selected for 5-7 days in the relevant antibiotic (puromycin 4 μg/ml; blasticidin 4 μg/ml). The selected cells were used as polyclonal populations to avoid possible biases resulting from the isolation of random clones. To express the luciferase gene in wild-type MC38 cells to facilitate tumor tracking (IVIS), we used the pLenti CMV V5-LUC Blast (w567-1) vector (Dr. Eric Campeau; Addgene Ref. 21474). Lentiviral transductions and cellular selection were carried out as described above.

Transgene expression was carried out via retroviral transductions using the P12-MMP retroviral vector to clone the relevant constructs, sometimes including an IRES-resistant cassette for subsequent selection. ATG16L1-depleted cells were reconstituted with FL or Nt (residues 1-299) forms of human ATG16L1, which includes two nucleotide differences in the sgRNA sequence that targets the endogenous mouse ATG16L1 form, but still retains the potential PAM. To ensure complete insensitivity of the human transgene to the action of the sgRNA directed to endogenous mouse ATG16L1, we introduced silent mutations in the target sequence of human ATG16L1 via site-directed mutagenesis using the mutagenic oligonucleotide: 5’ gcctccgcgccgctgacttcccTcgGtggaaAcg**<u>c</u>**acatc**<u>t</u>**cggagcaac 3’, where the capital letters indicate the mutations introduced: the first T mutates the PAM, the next G and A introduce additional silent mutations to cause more sequence divergence with endogenous mouse ATG16L1, and **<u>c</u>** and **t** highlight the nucleotides that are naturally different from the mouse cDNA. Mutant ATG16L1 FL and Nt (1-299) were cloned into the P12-MMP retroviral vector containing an IRES-hygromycin cassette for selection in 250 μg/ml hygromycin B. The mouse CD39-flag construct was ordered to VectorBuilder and cloned into a P12-MMP derivative including an IRES-blasticidin cassette for selection in 4 μg/ml blasticidin. GFP-LC3 and Cherry-LC3 fusion constructs were also subcloned into a P12-MMP IRES-blasticidin vector for selection as mentioned above. The DNA construct encoding the fusion protein GFP-SopF was kindly supplied by Dr. Felix Randow (LMB, Cambridge, UK) and subcloned into P12-MMP, in this case without a resistance cassette. Virus-containing supernatants were generated by co-transfecting HEK-293T cells with the relevant P12-MMP constructs and helper plasmids expressing gag-pol (pMD.gag-pol) and env (VSV-G; pMD-G). Retroviral transductions were carried out as described above for lentiviruses. All DNA constructs were verified by sequencing.

### Cell lysis and Western blot

To generate whole cell lysates for Western blotting, cell pellets were resuspended in 2x SDS sample buffer (SB) lacking β-mercaptoethanol and bromophenol blue (100 mM Tris-HCl pH 6.8, 4% SDS and 20% de glycerol) containing PMSF (1 mM; Merck, 10837091001) and a Protease Inhibitor Cocktail (PIC, 1/100, P8340, Sigma), boiled for 15 min and centrifuged (16100xg, 5 min, 4°C) to eliminate debris. Protein concentrations were measured using the DC Protein Assay Kit (Lowry method, BioRad, 5000116). Equal amounts of protein were diluted 1:1 with 2x SDS reducing sample buffer (2x RSB; SB containing β-mercaptoethanol (10 %) and bromophenol blue (0,3 %)), boiled and subjected to SDS-PAGE. Gels were then transferred to a polyvinilidene difluoride membrane (PVDF, Millipore, IPVH00010), and probed with primary antibodies against: ATG16L1 (mouse, MBL, M150-3, 1/1000), p62 (rabbit, MBL, PM045B, 1/1000), pMLKL (rabbit, Cell Signaling, 37333, 1/1000), MLKL (rabbit, Cell Signaling, 37705, 1/1000), cleaved caspase-3 (tCasp-3, rabbit, Cell Signaling, 9661, 1/1000), caspase 3 (rabbit, Cell Signaling, 14220, 1/1000), cytochrome c (rabbit, Proteintech, 10993-1-AP, 1/1000), LC3 (mouse, MBL, M186-3, 1/1000), GFP (mouse, Covance, MMS-118P, 1/5000), pSTING (rabbit, Cell Signaling, 72971S, 1/500), STING (rabbit, Proteintech, 19851-1-AP, 1/1000), BAK (rabbit, Cell Signaling, 12105, 1/1000), BAX (rabbit, Santa Cruz Biotech, N20, sc-493), Flag (rabbit, Cell Signaling, 14793, 1/1000), Actin (mouse, Sigma AC40, A3853, 1/1000) and GAPDH (mouse, Thermo Fisher, 6C5, AM4300, 1/1000). Secondary antibodies were purchased from Jackson Immunoresearch (goat anti-mouse HRP, 115-035-003, 1/10000; goat anti-rabbit HRP, 111-035-003, 1/10000). Blots were developed by chemiluminescence (ECL, Amersham, RPN2134). When required due to a molecular weight conflict, membranes were stripped in a 7 M guanidinium hydrochloride solution (10 min, RT), or the same samples rerun on a different gel using the same measurements of protein concentrations, for reprobing. Uncropped images of all Western blots are included in a Supplemental Material file.

### Cytochrome c release

Cells were lysed in a buffer containing 25 mM Tris-HCl pH 7.4, 10 mM KCl, 5 mM MgCl2, 1 mM EDTA, 250 mM sucrose and 0.02% digitonin for 2 min (ice), and centrifuged at 14000xg (4°C, 3 min). The resulting supernatants were collected and the corresponding pellets lysed in SB buffer. Samples were measured for protein concentration and Western blotting.

### Measurement of cell death

Both floating and attached cells were collected, centrifuged for 10 min (3300xg, 4°C) and resuspended in medium containing 1 μg/ml propidium iodide (PI) for A+S or DAPI (1 μg/ml) for samples containing MTX (since this drug emits red fluorescence). Cells were incubated for 10 min and then diluted 1:5 in medium before flow cytometry using a FACSCanto II device (Becton Dickinson) running the FACSDiva software (version 6.13). Cells were gated in the FS/SS window to include both live and dead cells (and exclude small debris) and assessed for PI or DAPI fluorescence (FL3). The percentage of PI/DAPI-positive cells (FL3) was established using the FlowJo (version 10) software (Becton Dickinson). Gating strategies are shown in Suppl. Fig. 5.

### Measurement of secreted ATP

Cells were seeded in 24-well plates and treated the next day. Around 5 h after treatment, 3 μM of the ecto-ATPase inhibitor ARL67156 (Sigma, A265) was added to the cultures. At the relevant experimental time points, supernatants were collected for ATP measurement using the Enliten ATP assay kit (Promega, FF2000) following the manufacturers’ instructions. To exclude possible variations in ATP secretion due to slightly different death kinetics, supernatants were harvested at two different times during death process: an early time point when cell death is around 40-50% and a late time point with cell death approaches 80-90%. These conditions were, respectively, 14 h and 20 h for MTX+zVAD, 16 h and 22 h for MTX, and 4h and 9 h for A+S. Cell death assays were carried out in parallel to verify the desired level of death at both time points as well as equal induction in FL and Nt cells. ATP secretion graphs display the cumulative data obtained from the sum of the luciferase activity units obtained at early and late times.

### Measurement of HMGB1 liberation by ELISA

Cells were plated in 24-well plates and treated with the different stimuli for the required times to reach around 70% cell death. At these time points, culture supernatants were collected, spun to remove cellular debris (10 min, 4°C, 3200xg) and tested for HMGB1 levels using an anti-mouse specific HMGB1 ELISA system (Invitrogen, Ref. EEL102) following the instructions provided by the manufacturers. HMGB1 concentrations were calculated using a standard curve generated as indicated in the product instructions. Cell death was measured in parallel to verify the desired level of death and equal induction in FL and Nt cells.

### Flow cytometry for CD39 and calreticulin

To establish the surface expression levels of CD39, cells were detached using Accutase (Sigma, A6964), resuspended in PBS and stained for 10 min on ice with Ghost Dye (Tonbo Biosciences, Ref. SKU-13-0865-T100; 1/1000) to discriminate live from dead cells. Cells were then washed, resuspended in PBS containing 2% BSA and 0.1% azide (FACS buffer) and stained with a primary anti-mouse CD39 antibody (rat, Biolegend, 143801, 1/1000) followed by a secondary anti-rat IgG Alexa Fluor 488 antibody (donkey, Thermo Fisher, A-21208, 1/500), in both cases for 40 min on ice with occasional mixing. Cells were then washed, fixed in 2% paraformaldehyde at room temperature, centrifuged again, resuspended in FACS buffer and analyzed in a FACSCanto II (Becton Dickinson) flow cytometer running the FACSDiva software (version 6.13). Subsequent analyses were done using FlowJo program (version 10). The gating strategy is shown in Suppl. Fig. 6A.

To assess the surface exposure of calreticulin during cell death, cells were treated in 24-well plates until they reached a level of cell death around 40-50%. Both floating and adherent cells were then collected from each well, in the latter case by detaching them using Accutase. Cells were resuspended in PBS and stained with Ghost Dye as described above. After washing, cells were resuspended in FACS buffer and stained with a primary anti-calreticulin antibody (rabbit, Cell Signaling, 12238, 1/1200) followed by a secondary anti-rabbit IgG Alexa Fluor 488 antibody (donkey, Thermo Fisher, A-21206, 1/400). The staining procedures, fixation and analysis by flow cytometry were done as described above for CD39. Measurements of cell death were carried out in parallel to verify the desired level of death as well as equal induction in FL and Nt cells. The gating strategy is shown in Supplementary Fig. 6B.

### Confocal microscopy

For immunofluorescence studies involving cells expressing GFP-LC3, cells were seeded on coverslips and treated the next day with the relevant stimuli. At the end of the assay, cells were washed with PBS and fixed in 4% paraformaldehyde (PFA) for 10 min. Samples were quenched/permeabilized in PBS containing 100 mM glycine and 0.5% Igepal CA-630 (RT, 30 min), incubated with DAPI (1 μg/ml, 5 min) and mounted in Prolong Gold anti-fade medium (Thermo Fisher, P10144). Pictures were taken in a Stellaris 8 STED confocal microscope (Leica; LAS X 4.2.0.23645 software) using the 488 nm (GFP-LC3; detection window 505-545 nm) and 405 nm (DAPI; detection 415-475 nm) laser lines. The blue signal represents DAPI-stained nuclei in all pictures. Scale bars represent 10 μm in all micrographs.

To evaluate vesicular ATP content cells expressing Cherry-LC3 were seeded on coverslips, treated the next day and, at the end of the experiment, stained with quinacrine or Bodipy-ATP. For quinacrine staining, cells were washed with Krebs-Ringer buffer (25 mM Hepes pH 7.4; 125 mM NaCl; 5 mM KCl; 1 mM MgSO_4_; 0.7 mM KH_2_PO_4_; 2 mM CaCl_2_; 6 mM glucose) and incubated with the same buffer containing 5 μM quinacrine for 15 min at 37°C. For staining with Bodipy-ATP, cells were incubated in DMEM culture medium containing 40 μM Bodipy-ATP for 3 h at 37°C. Coverslips were washed in the respective buffers, placed upside down on glass-bottom Wilco plates (GWSt-3522) and analyzed *in vivo* on an Okolab chamber at 37°C and 5% CO_2_ coupled to the same Stellaris 8 STED confocal microscope (Leica) mentioned above, using in this case the 488 nm (for quinacrine and Bodipy-ATP, detection window 500-550 nm), and 555 nm (for Cherry-LC3, detection window 563-630 nm) laser lines.

Quantitation of vesicle number was done in all cases by blindly counting vesicles on confocal micrographs. Colocalizations were quantified by measuring the Manders coefficient using the JACoP plugin of the Fiji program for image analysis. Scale bars represent 10 μm in all micrographs. All experiments were repeated at least three times with qualitatively similar results.

### Tumor vaccination assays and evaluation of tumor growth

Female C57BL/6 mice (5-7 weeks old) were obtained from Envigo. Mice were vaccinated with 5.5×10^6^ MC38 cells treated with MTX (1.5 μM) + zVAD.fmk (20 μM) for 15 h, conditions that produced around 50% cell death. Floating cells were collected and pooled together with the remaining attached cells after detachment with trypsin (except for the experiments involving CD39-expressing cells, where Accutase was used instead). Cells were washed with PBS, resuspended in 200 μl of PBS containing ARL67156 (30 μM) and injected subcutaneously in the left flank. One week later mice were challenged in the right flank with wild-type MC38 cells stably transduced with a vector constitutively expressing the firefly luciferase gene (1.35×10^5^ cells per mice in 200 PBS μl). Initial control experiments showed no tumor growth after vaccination, nor post-vaccination tumors ever grew in the left flank. Tumor volume was measured with a digital caliper at days 6, 10 and 14-15 after challenge with MC38 cells. Volume was calculated with the formula V=(W^2^xL)/2, where V is volume, W is width and L is length. Body weight and general health status were monitored daily during the tumor growth period. At days 14-16 post-inoculation some of the tumors generated in the most permissive conditions usually reached the humaine endpoint established at 1500 mm^3^ of tumor volume and required euthanization. Accordingly, experiments were normally limited to a 14-15 days of tumor growth. Mice were sacrificed by carbon dioxide inhalation at the end of the assay or after reaching the humane endpoint criteria.

To evaluate tumor bioluminescence activity^67^, mice were intraperitoneally inoculated with 200 μl per mice of a solution of D-luciferin potassium salt (Revvity, Ref. 122799; 15 mg/ml in PBS; 150 mg/kg) as a luciferase substrate. Mice were then anesthetized via isoflurane inhalation (4% induction, 2% maintenance) and analyzed using an IVIS device (IVIS Lumina II, Caliper Life Sciences) running on a Living Image 4.8.2 software. A series of bioluminescence intensity images were sequentially acquired and subsequently analyzed to select those showing maximum activity for each mouse and measurement. Image quantification was carried out by assessing the total flux (photons per second) provided by a common ROI that includes the bioluminescent area obtained for all mice analyzed in a single experiment. IVIS measurements normally required higher n to reach significance compared to caliper assessments.

Depletion of T cells was achieved by repeated intraperitoneal injection of an anti-Thy1.2 antibody (anti-Thy1.2, Leinco, C393) or an isotype control antibody (control rat IgG, Leinco, I-1034). Mice received an initial 900 μg dose (diluted in 200 μl PBS) three days after vaccination (four days before challenge), followed by repeated 300 μg maintenance doses every 3 days until the end of the experiment. Tumor growth was evaluated over time as described above. Depletion was judged at the end of the assay by measuring the percentage of CD3-positive cells in spleen samples by flow cytometry. Spleens were extracted and mechanically disrupted against a 70 μm cell strainer using a syringe plunger. Cells were recovered in RPMI culture medium containing 10% FCS. Red blood cells were then lysed using ACK buffer. The remaining cellular suspension was stained with Ghost Dye 780, incubated with an Fc block reagent, stained for CD3 and analyzed by flow cytometry using the same reagents, procedures, flow cytometer and software tools as described for evaluation of the immune profile of MC38 tumors. The gating strategy is displayed in Suppl. Fig. 7.

### Evaluation of the immune profile of MC38 tumors by flow cytometry

At the end of the experiment tumors were dissected, placed in RPMI culture medium supplemented with 10% FCS and processed roughly following a previously established protocol^68^. Comparable tumor portions were then excised, minced and digested in RPMI without FCS containing 0.2 mg/ml Liberase TM (Sigma, Ref. 5401119001) and 240 μg/ml (DNAse I, Ref. 11284932001) for 30 min at 37°C with gentle shaking. Samples were then filtered with a 70 μm cell strainer (Falcon, 352350), centrifuged (500xg, 10 min, 4°C) and resuspended in ACK buffer (Thermo Fisher A1049201) for erythrocyte lysis (3 min, RT). Cells were counted and 10^6^ cells per staining spun, resuspended in PBS and stained with Ghost Dye 780 (5 min, ice, 1/4000) to detect dead cells. Samples were then washed, resuspended in FACS buffer (PBS, 2% BSA, 0.1% azide) and incubated with an Fc blocking reagent (TruStain FcX, anti-mouse CD16/32, Biolegend, 156603, 1/300) for 30 min on ice. Cells were washed again and stained separately for the different lymphoid and myeloid markers. Lymphoid antibodies were: anti-CD45 PE/Cyanine7 (Biolegend, 109829, 1/400), anti-CD3 Brilliant Violet 421 (Biolegend, 100227, 1/100), anti-NK1.1 PE/Dazzle 594 (Biolegend, 108747, 1/150), anti-CD4 FITC (Biolegend, 100405, 1/200), anti-CD8a PE/Cyanine5 (Biolegend, 100709, 1/500), anti-CD25 Pacific Blue (Biolegend, 102021, 1/75), anti-CD69 Brilliant Violet 650 (Biolegend, 104541, 1/200). Myeloid antibodies: anti-CD45 PE/Cyanine7 (Biolegend, 109829, 1/400), anti-CD11b BB515 (Becton Dickinson, 564454, 1/400), anti-CD11c Brilliant Violet 421 (Biolegend, 117329, 1/50), anti-F4/80 Alexa Fluor 647 (Becton Dickinson, 565853, 1/200). Samples were then washed, fixed in 2% paraformaldehyde and run on a Cytek Aurora 4L flow cytometer using the SpectroFlo v3 program. Subsequent analyses were done using the FlowJo (version 10) software. The gating strategies followed for both lymphoid and myeloid samples are shown in Suppl. Figs. 8A and 8B, respectively.

### Animal experimentation

Wild-type C57BL6/J mice used for the tumor vaccination studies were hosted in a specific pathogen-free facility including ventilated racks with controlled temperature (23°C), humidity (50%) and light/dark cycles (12 h/12 h), and access to food and water *ad libitum*. Animal care protocols followed the ARRIVE (https://arriveguidelines.org/) guidelines, in strict accordance with the European Commission legislation for the protection of animal use for experimental purposes (2010/63/EU). All animal procedures were approved by the Ethics and Animal Wellbeing committees of the research institute (Centro de Biología Molecular Severo Ochoa, CBMSO), the hosting institution (Consejo Superior de Investigaciones Científicas, CSIC), and the Dirección General del Medio Ambiente of the Comunidad de Madrid local government.

## Statistics

Graphs displaying cell death (flow cytometry), ATP secretion (luciferase activity), HMGB1 liberation (ELISA) and calreticulin exposure (flow cytometry) show means -/+ standard deviations obtained from n=3 experimental points (n; biological replicas). Results including vesicle counts display around 25 data points and are presented as box-plots where the central line represents the median value, the box shows percentiles 25-75 and the whiskers include the most extreme values. In both cases, statistical significance (*P*-values) was calculated using two-tailed Student’s *t*-tests. Flow cytometry data displaying tumor immune profiles show mean values -/+ standard deviations obtained from n=18 experimental points. Statistical significance in this case was calculated using a Mann-Whitney non-parametric t test. Graphs showing tumor growth over time display mean values + standard deviations of a variable number of experimental replicas (n varies between 5 and 22, as indicated in the respective figure legends), and statistical significance was calculated using a two-way ANOVA with Tukey’s multiple comparison test. Graphs and statistical analyses were obtained using GraphPad Prism 9. The displayed results are representative of at least 3 repetitions.

## Data availability statement

Data and materials are freely available upon request to the corresponding author.

## Acknowledgements

We thank Rocío Díaz and Tamara Rosell for experimental help. We also thank the CBM Flow Cytometry, Advanced Microscopy, Animal House and Preclinical Biomedicine facilities and their personnel for support. This work was funded by grants PID2020-114699RB-I00 and PID2023-148747OB-I00 from the Ministerio de Ciencia y Tecnología (Spanish Government; MCIN/AEI/10.13039/501100011033, FEDER, UE), IDEAS18093PIME from the Asociación Española Contra el Cáncer (AECC) and SA042P17 from the Junta de Castilla y León local government. Additional funding comes from the of Excellence Severo Ochoa Grant CEX2021-001154-S from MCIN/AEI/10.13039/501100011033. J.B.L is the recipient of a predoctoral fellowship from the Formación de Personal Investigador (FPI) program of the Spanish Ministry of Science and Technology (Ref. PRE2021-097461; MCIU/AEI/10.13039/501100011033; Spanish Government and FSE+). D.O.S. is the recipient of a Predoctoral Fellowship from the Comunidad de Madrid local government (Ref. PIPF-2023_SAL-GL-29621). The CBM receives institutional support by Fundación Ramon Areces. F.X.P. holds a tenured position at CSIC.

## Author contributions

J.B.L. and D.O.S. performed the experiments; K. S. provided experimental support. F.X.P. conceived and designed the study, and wrote the manuscript.

## Supplementary Figures and Figure Legends

**Supplementary Figure 1.**
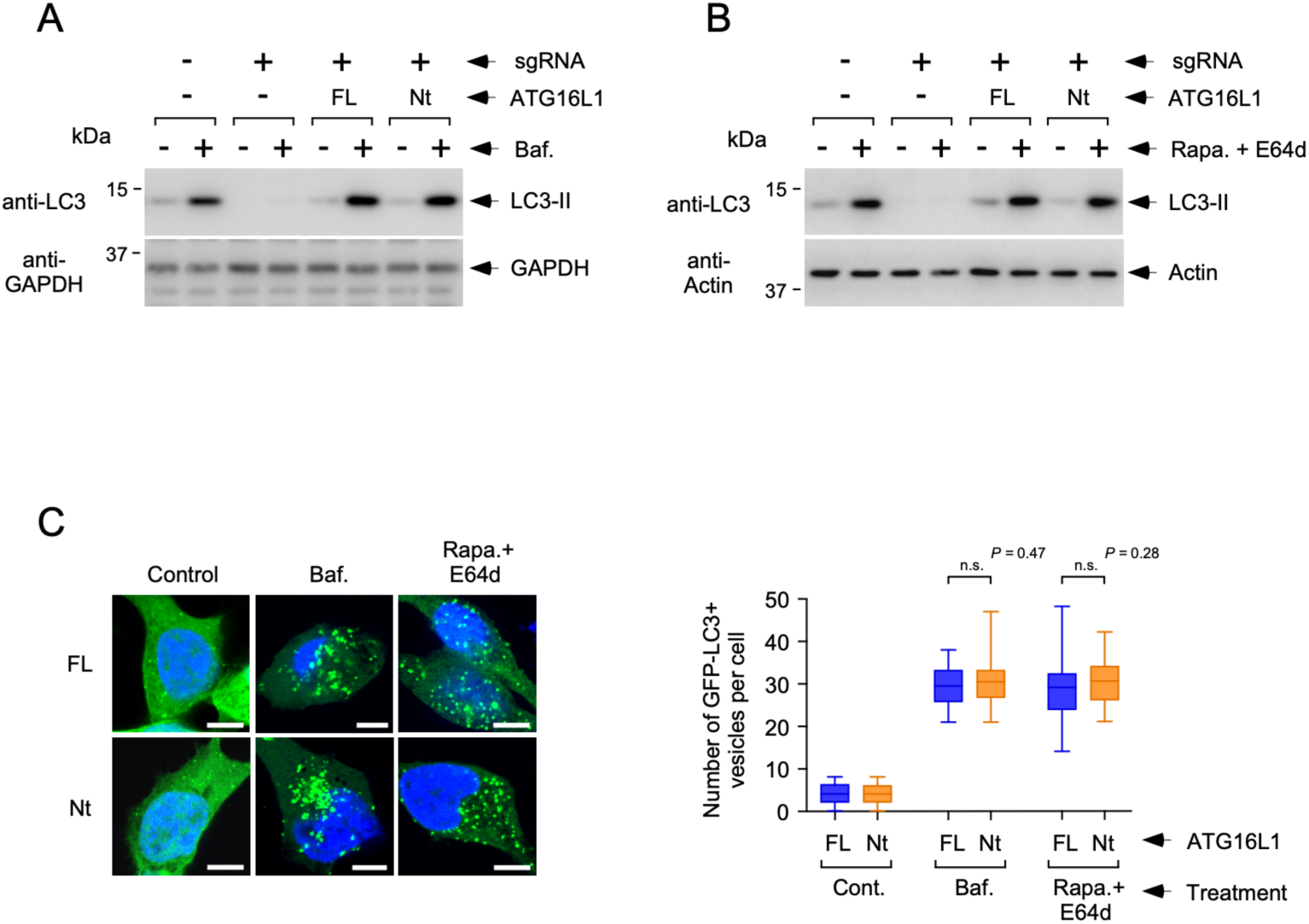
Comparable levels of canonical autophagy in engineered MC38 cells expressing FL and Nt forms of ATG16L1. **(A)** The indicated MC38 strains were treated with bafilomycin (Baf., 50 nM) for 8 h and lysed for Western blot against the shown molecules. **(B)** The indicated MC38 strains were treated with rapamycin (Rapa., 2 μM) and E64d (10 μg/ml) for 8 h and lysed for Western blot against the shown molecules. **(C)** The indicated MC38 strains expressing GFP-LC3 were treated as in A and B (as displayed) and fixed for confocal microscopy. Shown are representative confocal pictures (left panel) and quantification of the number of GFP-LC3-puncta per cell (right panel). Data are displayed as box-plots where the central line indicates the median value, the box shows percentiles 25-75 and the whiskers include the most extreme values (n=26 cells; n.s. *P*>0.05 two-tailed Student’s t test).

**Supplementary Figure 2.**
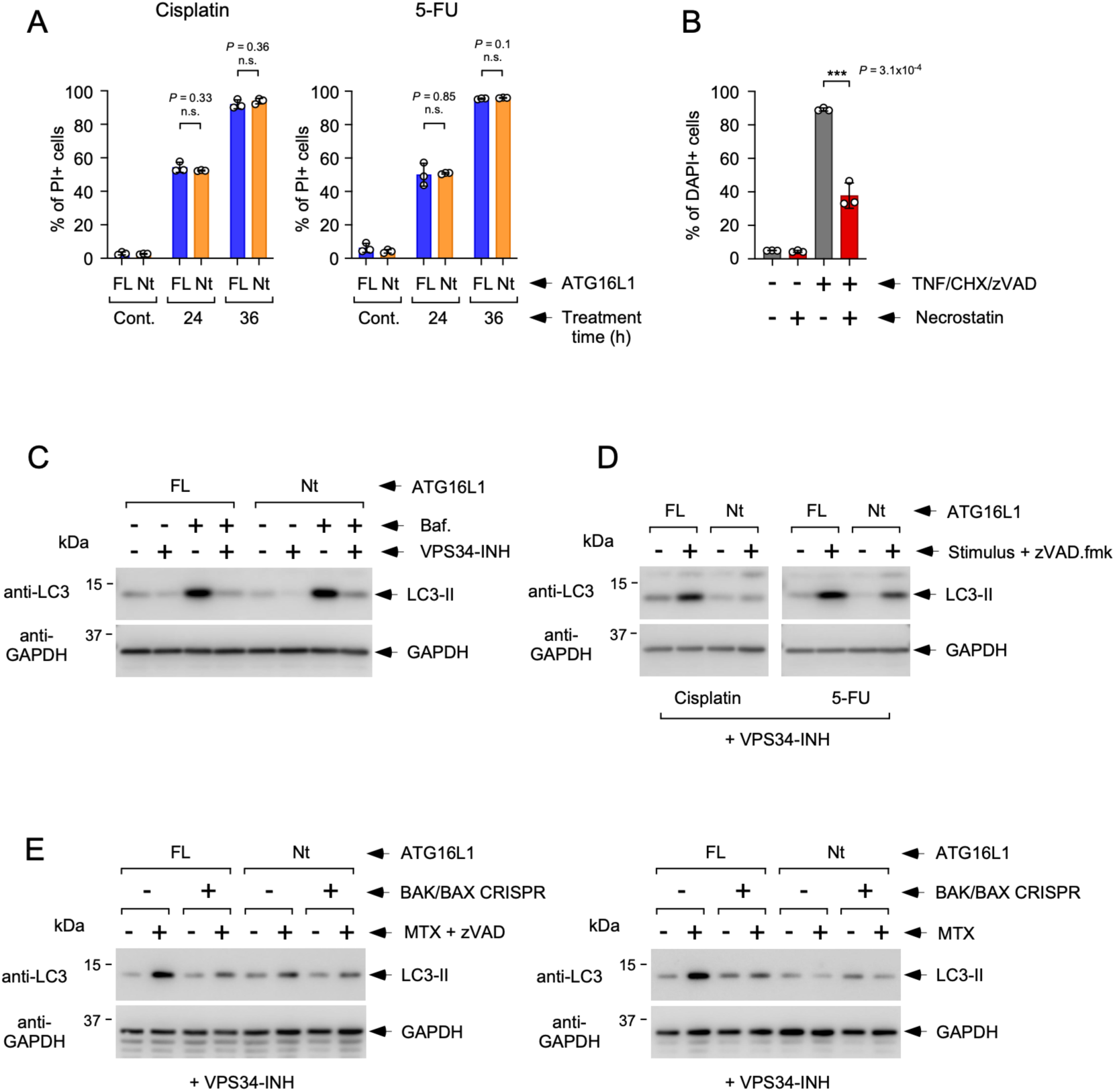
Cell death and autophagic response of MC38 cells expressing ATG16L1 FL or Nt treated with different stimuli. **(A)**. Cell death response to cisplatin and 5-FU. The displayed cellular strains (FL or Nt) were treated for the indicated times with cisplatin (25 μM, left panel) or 5-FU (150 μM, right panel), and stained with propidium iodide (PI) to measure cell death by flow cytometry. Shown are mean values -/+ s.d. of the percentage of PI-positive cells obtained from triplicate experimental points (n=3 biological replicas; n.s. *P*>0.05 two-tailed Student’s t test). **(B)** Inhibition of necroptosis in MC38 cells by necrostatin. MC38 cells engineered to express ATG16L1-FL were treated with TNF (20 ng/ml), cycloheximide (CHX, 10 μg/ml) and zVAD.fmk (20 μM) for 18 h in the absence or presence of necrostatin (25 μM, as indicated) and stained with DAPI to measure cell death by flow cytometry. Shown are mean values -/+ s.d. of the percentage of DAPI-positive cells obtained from triplicate experimental points (n=3 biological replicas; \*\*\**P*<0.001 two-tailed Student’s t test). **(C)** Inhibition of basal autophagy by VPS34-INH in MC38 cells. MC38 FL and Nt strains were treated with bafilomycin (50 nM, 8 h) in the absence or presence of VPS34-INH (2 μM) as indicated, and lysed for Western blot against the shown molecules. **(D)** Unconventional autophagy induced by cisplatin and 5-FU. The indicated cells were treated with cisplatin (25 μM, 20 h, left panel) or 5-FU (150 μM, 20 h, right panel) and zVAD.fmk (20 μM), in the presence of VPS34-INH (2 μM), and lysed for Western blot against the shown molecules. **(E)** FL or Nt cells further manipulated to deplete BAK and BAX (as indicated; see Fig. 2C) were treated for 16 h with MTX+zVAD (1.5 μM and 20 μM, respectively, left panel) or for 18 h with MTX (1.5 μM, right panel), in the presence of VPS-34-INH (2 μM), and lysed for Western blot using the indicated antibodies.

**Supplementary Figure 3.**
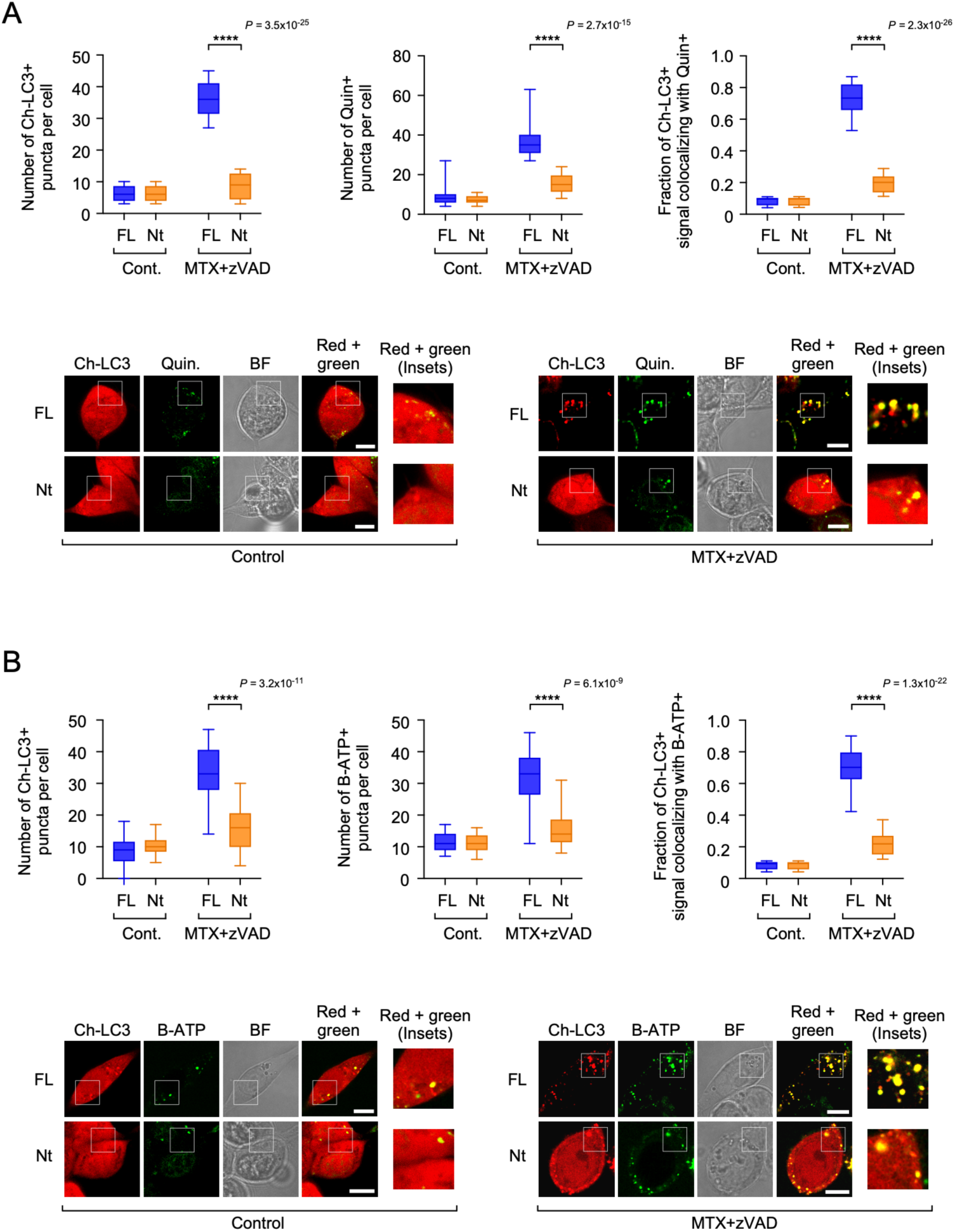
The LC3-positive vesicles generated in response to MTX+zVAD are stained with the ATP-sensitive dyes quinacrine and Bodipy-ATP. (A-B) The indicated FL and Nt cells expressing Cherry-LC3 (Ch-LC3) were treated for 13 h with MTX+zVAD (1.5 μM and 20 μM, respectively) in the presence of VPS34-INH (2 μM), stained with quinacrine (Quin.; **A**) or Bodipy-ATP (B-ATP, **B**) and analyzed *in vivo* by confocal microscopy. The graphs on the top panels show the number of Cherry-LC3-positive vesicles per cell (left), the number of quinacrine **(A)** or Bodipy-ATP **(B)**-positive vesicles per cell (middle) and the fraction of Cherry-LC3-positive signal that colocalizes with the green signal (right). Data are presented as box-plots where the central line represents the median value, the box shows percentiles 25-75 and the whiskers include the most extreme values. Statistical significance (*P*-values) was calculated using two-tailed Student’s *t*-tests (\*\*\*\**P*<0.0001). The bottom panels show representative confocal pictures (BF, bright field).

**Supplementary Figure 4.**
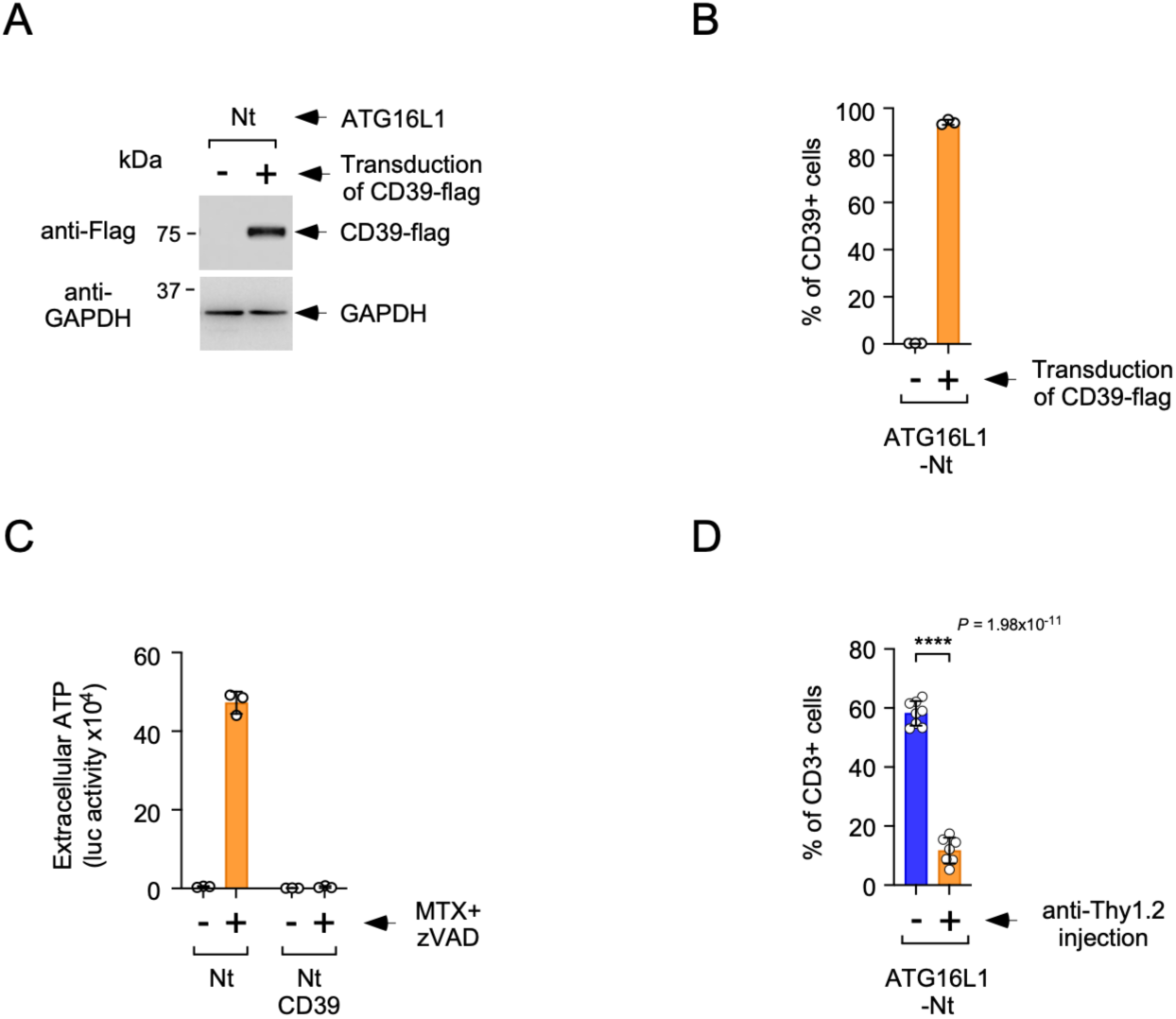
Evaluation of CD39 expression and function in MC38-Nt cells transduced with a retrovirus carrying a mouse CD39-Flag construct, and evaluation of T cell depletion in mice injected with anti-Thy1.2 antibodies. **(A)** MC38 cells expressing ATG16L1-Nt were transduced with a retroviral construct expressing CD39-Flag (+) or a control construct (-), selected in blasticidin and lysed for Western blot against the indicated molecules. **(B)** Surface expression levels of CD39. The cells described in A were stained with an anti-CD39 antibody and subjected to flow cytometry. The graph shows mean values -/+ s.d. of the percentage of CD39-positive cells (n=3 biological replicas). **(C)** CD39 expression reduces the levels of extracellular ATP during cell death. The indicated cells were treated with MTX+zVAD (1.5 μM and 20 μM, respectively) and the levels of extracellular ATP were measured at 14 and 20 h time points. The graphs show mean values -/+ s.d. of cumulative data resulting from the sum of luciferase activity units obtained at both time points from triplicate experimental points (n=3 biological replicas). **(D)** Depletion of T cells by injection of anti-Thy1 antibodies. Spleen cells were obtained from mice subjected to recurrent injection of control (-) or anti-Thy1.2 (+) antibodies and stained for CD3 expression. The graph shows mean values-/+ s.d. of the percentage of CD3-positive cells in the indicated samples (n=8 for control mice and n=7 for the anti-Thy1.2 group; **** *P*<0.0001 two-tailed Student’s t test).

**Supplementary Figure 5.**
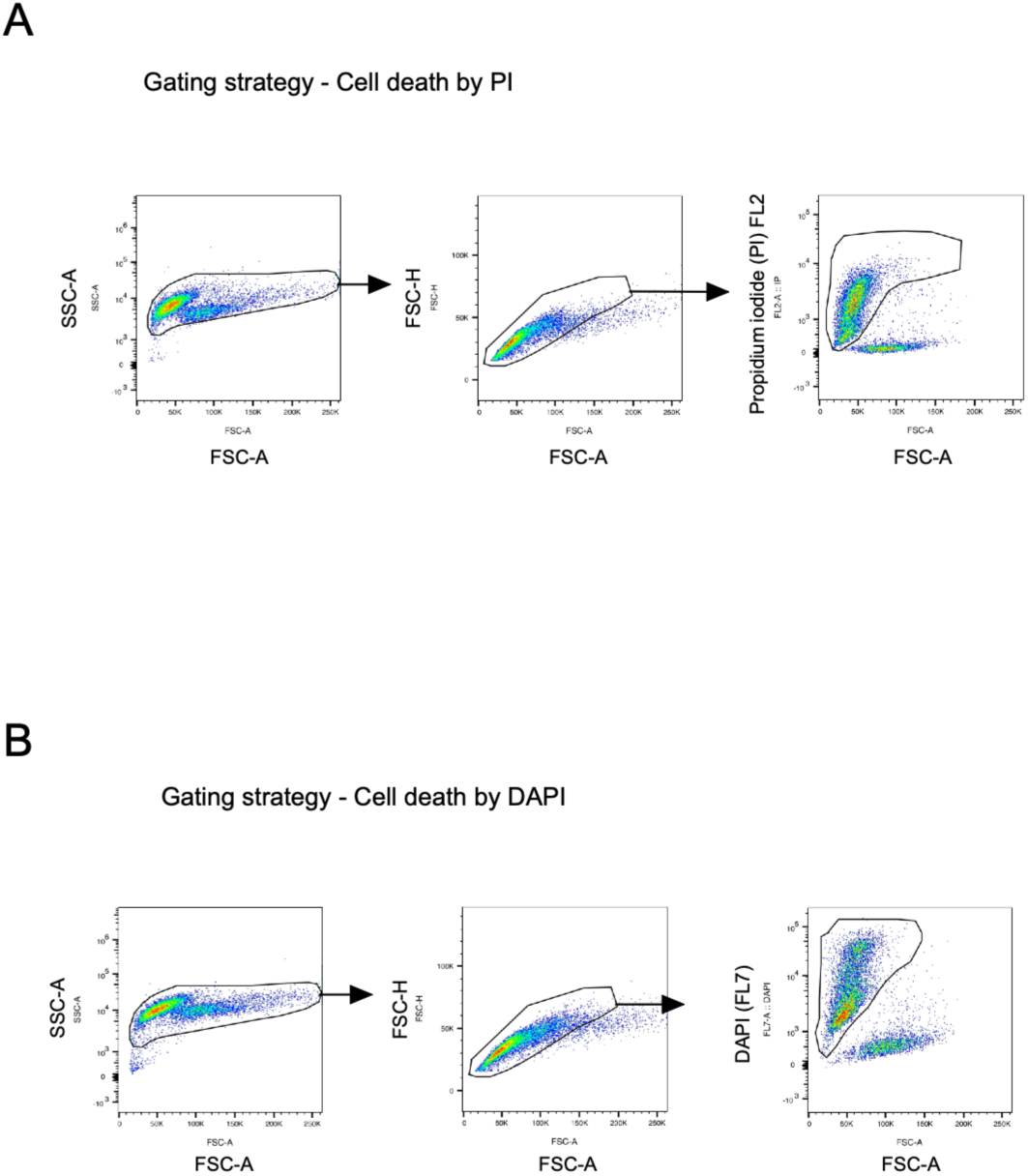
Gating strategies to assess cell death by flow cytometry. **(A)** Gating strategy used to measure cell death by propidium-iodide (PI) staining. **(B)** Gating strategy used to measure cell death by DAPI staining.

**Supplementary Figure 6.**
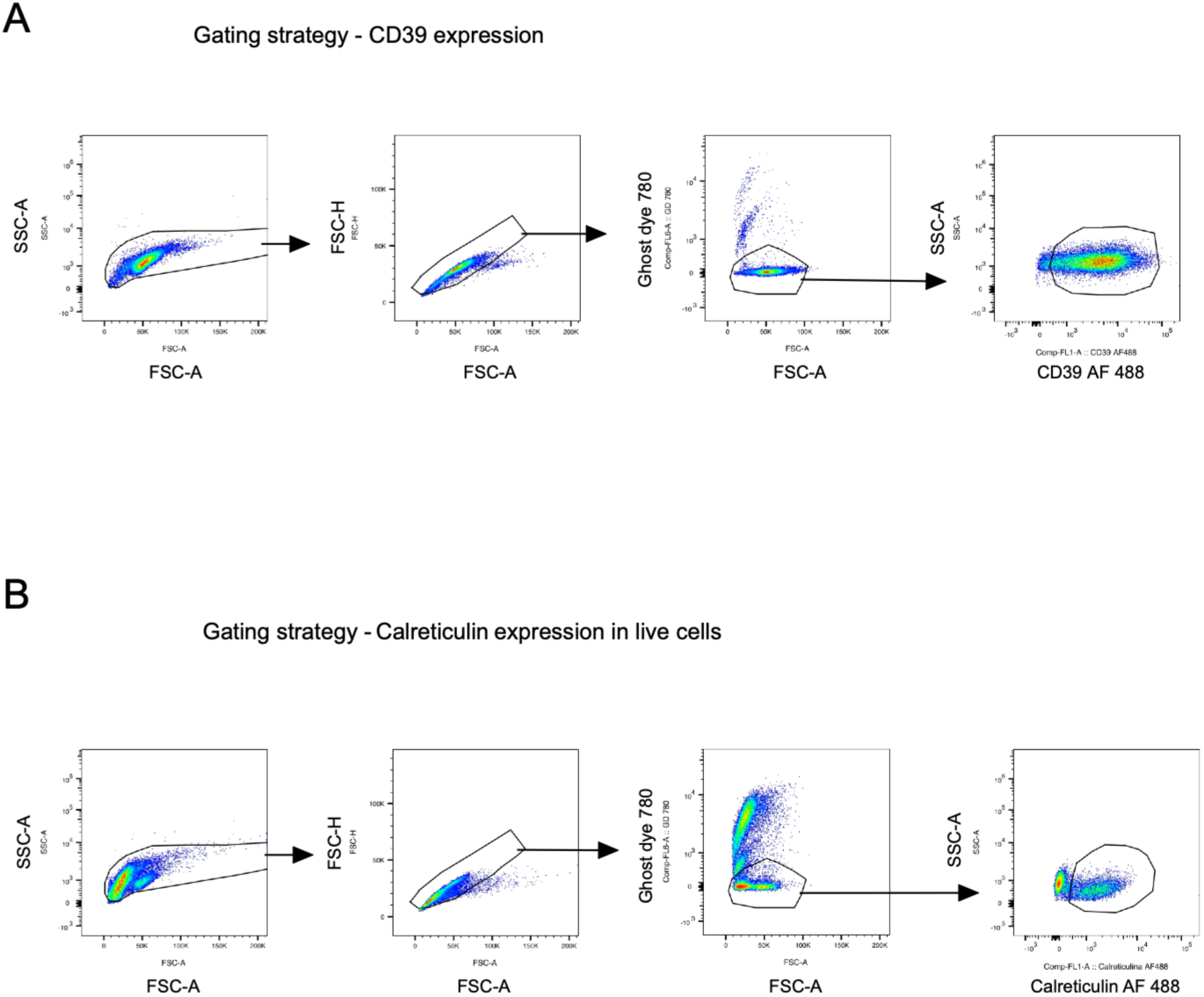
Gating strategies to assess CD39 and calreticulin plasma membrane expression levels by flow cytometry. **(A)** Gating strategy used to measure CD39 expression. **(B)** Gating strategy used to measure calreticulin expression in live cells.

**Supplementary Figure 7.**
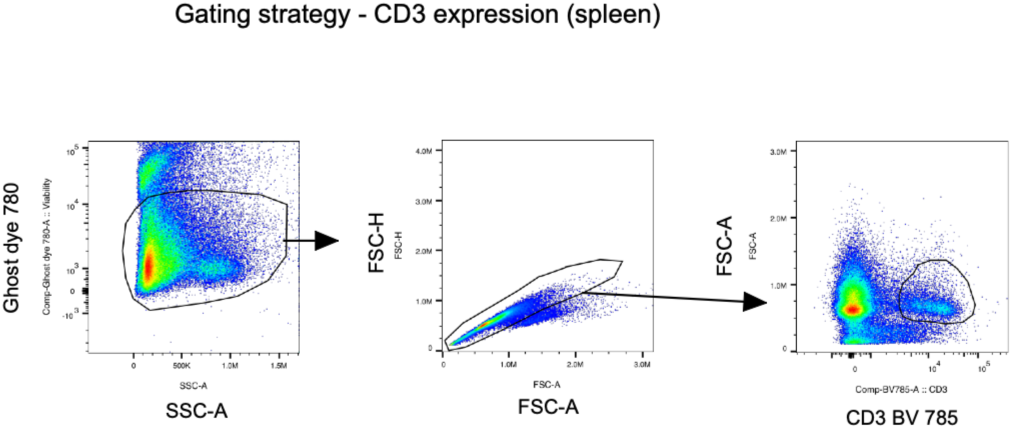
Gating strategy to measure T cell depletion caused by anti-Thy1.2 injection (spleen samples).

**Supplementary Figure 8.**
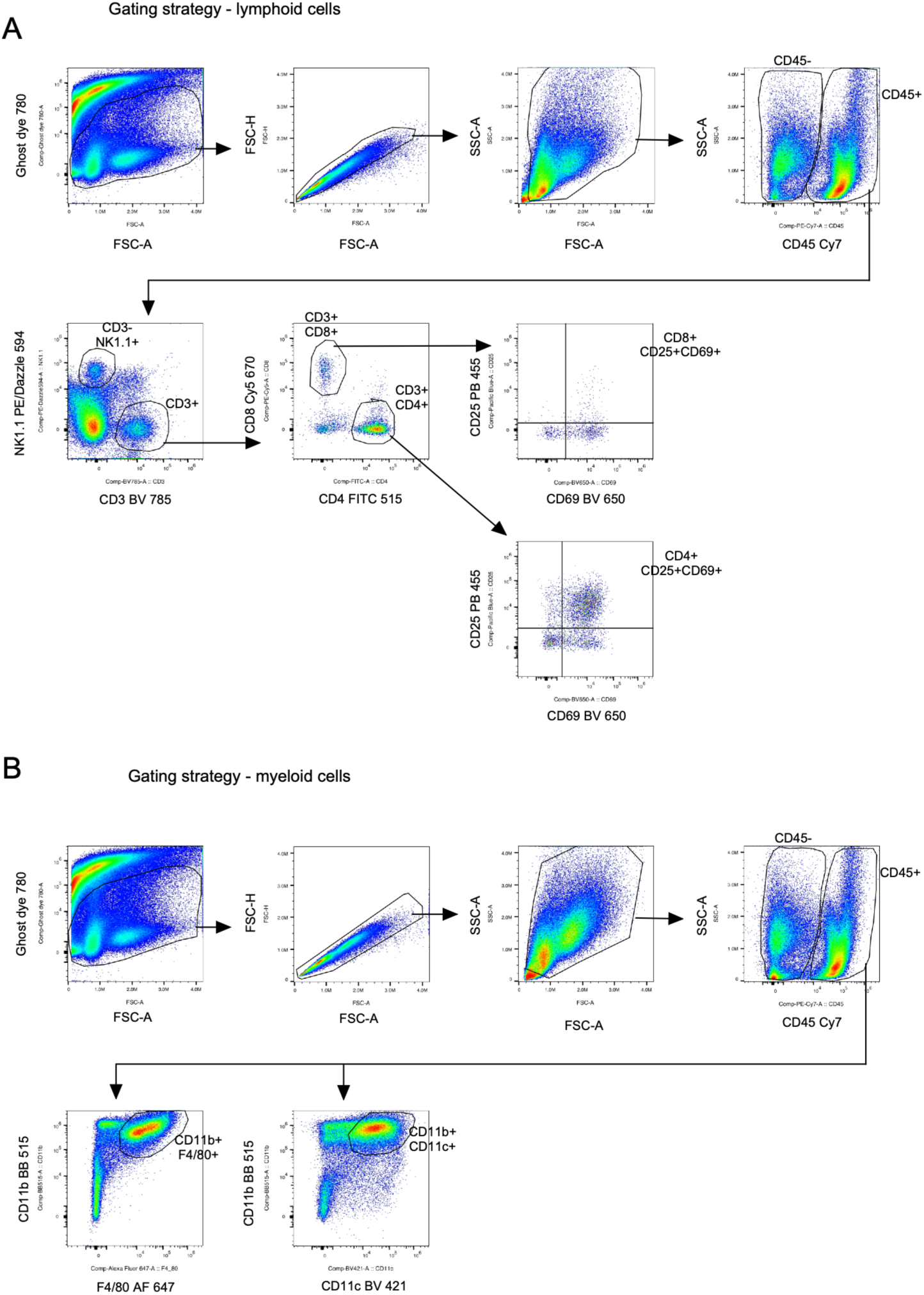
Gating strategies to assess the immune infiltration profiles of MC38 tumors. **(A)** Gating strategy for lymphoid populations. **(B)** Gating strategy for myeloid populations.

